# Alzheimer’s disease brain-derived tau-containing extracellular vesicles: Pathobiology and GABAergic neuronal transmission

**DOI:** 10.1101/2020.03.15.992719

**Authors:** Zhi Ruan, Dhruba Pathak, Srinidhi Venkatesan Kalavai, Asuka Yoshii-Kitahara, Satoshi Muraoka, Kayo Takamatsu-Yukawa, Nemil Bhatt, Jianqiao Hu, Yuzhi Wang, Samuel Hersh, Santhi Gorantla, Rakez Kayed, Howard E. Gendelman, Seiko Ikezu, Jennifer I. Luebke, Tsuneya Ikezu

## Abstract

Extracellular vesicles (EVs) propagate tau pathology for Alzheimer’s disease (AD). How EV transmission influences AD are, nonetheless, poorly understood. To these ends, the physicochemical and molecular structure-function relationships of human brain-derived EVs, from AD and prodromal AD (pAD), were compared to non-demented controls (CTRL). AD EVs were shown to be significantly enriched in epitope-specific tau oligomers versus pAD or CTRL EVs assayed by dot-blot and atomic force microscopy tests. AD EVs were efficiently internalized by murine cortical neurons and transferred tau with higher aggregation potency than pAD and CTRL EVs. Strikingly, inoculation of tau-containing AD EVs into the outer molecular layer of the dentate gyrus induced tau propagation throughout the hippocampus. This was seen in 22 months-old C57BL/6 mice at 4.5 months post-injection by semiquantitative brain-wide immunohistochemistry tests with multiple anti-phospho-tau (p-tau) antibodies. Inoculation of the equal amount of tau from CTRL EVs or as oligomer or fibril-enriched fraction from the same AD donor showed little propagation. AD EVs induced tau accumulation in the hippocampus as oligomers or sarkosyl-insoluble proteins. Unexpectedly, p-tau cells were mostly GAD67^+^ GABAergic neurons and to a lesser extent, GluR2/3^+^ excitatory mossy cells, showing preferential EV-mediated GABAergic neuronal tau propagation. Whole-cell patch clamp recording of Cornu Ammonis (CA1) pyramidal cells showed significant reduction in the amplitude of spontaneous inhibitory post-synaptic currents. This was accompanied by reductions in c-fos^+^ GAD67^+^GABAergic neurons and GAD67^+^ GABAergic neuronal puncta surrounding pyramidal neurons in the CA1 region confirming reduced interneuronal projections. Our study posits a novel tau-associated pathological mechanism for brain-derived EVs.

## Introduction

Accumulation of the misfolded microtubule-associated protein tau is a neuropathological hallmark of Alzheimer’s disease (AD). Tau is closely associated with AD cognitive decline [3]. Abnormally aggregated and phosphorylated tau (p-tau) first appears in the entorhinal cortex at the disease’ prodromal stage spreading in hierarchical patterns to the hippocampal regions then throughout the neocortex [9]. A growing body of evidence supports a prion-like cell-to-cell transmission for tau. Indeed, extracellular tau is seen to be internalized into healthy cells, where templated misfolding occurs leading to tau aggregates. This is then followed by another cycle of tau spread heralded by its cell-based secretion. Tau is mostly secreted in free form. A minor fraction of tau is associated with extracellular vesicles (EVs) in the cerebrospinal fluid (CSF) and blood of both AD and control (CTRL) patients [2, 10, 28, 61, 76]. The levels of free tau in CSF and in neuron-derived plasma EV tau from patients with mild cognitive impairment (MCI) or AD correlate with disease progression [2, 72] suggesting potential pathogenic roles of both forms of tau. Whether EV and free form tau contribute differently to tau propagation remains unresolved. A disease-associated role for paired helical filament (PHF)-tau from AD or other tauopathy brains was demonstrated following its inoculation into mouse brains leading to tau neuropathology [33, 54]. Notably, EVs isolated from transgenic tau mouse brains, AD plasma, or human induced pluripotent stem cells (iPSCs) expressing recombinant mutant tau also initiate propagation of tau in mouse brain tissues [5, 57, 71]. Pharmacologic inhibition of exosome synthesis significantly reduces tau propagation [4, 8]. The molecular mechanisms of cell-to-cell transmission of EV and free tau aggregates via uptake and secretion were subjects of intense investigation [10, 16, 60]. While the mode of uptake of free tau appears dependent on its conformational and post-translational modifications [24, 38, 48], EV tau uptake is affected by its surface proteins. EVs can target specific cell types by the interaction between EV and cell surface proteins [68]. To understand molecular composition of human brain-derived EVs, we have recently developed a separation protocol of human and mouse brain-derived EVs, by which we successfully enriched EVs with limited contamination from cytosolic components including the endoplasmic reticulum and Golgi [51, 53]. Our proteomic profiling of AD and CTRL brain-derived EVs identified glia-derived EV molecules enriched in AD cases and significantly differentially expressed proteins, which can distinguish AD from CTRL cases with 88% accuracy by a machine learning approach [51]. Furthermore, EVs isolated from interleukin (IL)-1β-stimulated human primary astrocytes showed increased expression of integrin-β3 (ITGB3), which was critical for neuronal EV uptake [74]. These data demonstrate that disease-associated pathologies such as glial inflammation can alter the molecular composition and neuronal uptake of EVs affecting their potency of tau spread.

There has been no comprehensive analysis of tau pathology development after the injection of human brain-derived EVs from CTRL or AD patients. Moreover, to fully understand the difference in potency between EV-associated and vesicle free tau, it is critical to compare propagation induced by different form of tau isolated from the same donor. Here we aimed to characterize brain-derived EVs separated from AD, prodromal AD (pAD) and age/sex-matched CTRL for their biophysical, biochemical, and neurobiological properties as well as for tau pathology after injection in the outer molecular layer (OML) of dentate gyrus (DG) in aged C57BL/6 (B6) mice. The recipient mice were tested by immunohistochemical and biochemical characterization of tau accumulation in the hippocampus. We also assessed the difference in tau pathology development after intrahippocampal injections of EV-, tau oligomer- and tau fibril-enriched fractions in mice. Finally, CA1 pyramidal neurons in the hippocampus of the recipient mice were assessed with whole-cell patch clamp recording to determine whether tau accumulation induces alterations in neurophysiological function.

## Results

### Detection of tau oligomers in AD and pAD brain-derived EVs

EVs consist of cell-derived lipid bilayer classified as exosomes or microvesicles. Exosomes are 30-150nm in size secreted after the fusion of endosomes with cell surface. Microvesicles are 100-1000nm in size secreted by outward budding of plasma membranes [21, 22, 60, 75]. They were originally part of the clearance system of unmetabolized cell composites. However, accumulative evidence suggests that EVs play critical roles for spreading pathological proteins. In this way they contribute to the pathobiology of neurodegenerative diseases [4, 18, 27, 32]. While tau is found both in exosomes and microvesicles from tauopathy mouse brains, neuroblastoma cells, CSF, and plasma in AD patients [2, 23, 28, 61] no study to date, has reported detailed analysis of AD brain-derived EVs. We first isolated exosome-enriched EV fractions from AD, pAD, and CTRL patient brain samples using the previously published protocol (Supplementary Table S1 for patients’ demographics) [51, 53]. Briefly, fresh-frozen post mortem brain tissues containing gray matter were lightly minced followed by the sequential centrifugation and discontinuous sucrose density gradient ultracentrifugation (Fig. 1a). Analyses of the isolated fractions by transmission electron microscopy (TEM, Fig. 1b) and nanoparticles tracking analysis (NTA, Fig. 1c) demonstrated enrichment of brain EVs with a size of exosomes [11]. There was no difference in terms of EV particle concentration or size among groups (Fig. 1d-e). ELISA analysis of total tau, Aβ40, and Aβ42 in EVs show total tau abundance without Aβ40; whereas Aβ42 are enriched in AD EVs (Supplementary Table S2). Considering that tau oligomers are in the nanometer size [17], we postulated that brain-derived EVs contain tau in oligomer forms. Indeed, there was a significantly higher amount of oligomeric tau in EVs derived from AD compared to CTRLs. These data were confirmed by tau oligomer-specific monoclonal antibodies TOMA-1 and TOMA-2, but not by TOMA-3 or TOMA-4 (Fig. 1f-I and Supplementary Fig. S1). There was no difference in immunoreactivity confirmed by tau oligomer polyclonal antibodies T22 and T18 among three groups (Fig. 1j-k). In addition, atomic force microscopy (AFM) analysis of detergent-insoluble fraction of AD and pAD but not of CRTL EVs detected globular particles. These were uncovered at the mode height 4-6 nm consistent with tau oligomers (Fig. 1l-m). Taken together, these data suggest that AD and pAD EVs are enriched in tau oligomers compared to CTRLs, indicating EV tau seeding potency and pathogenic activities.

**Figure 1.**
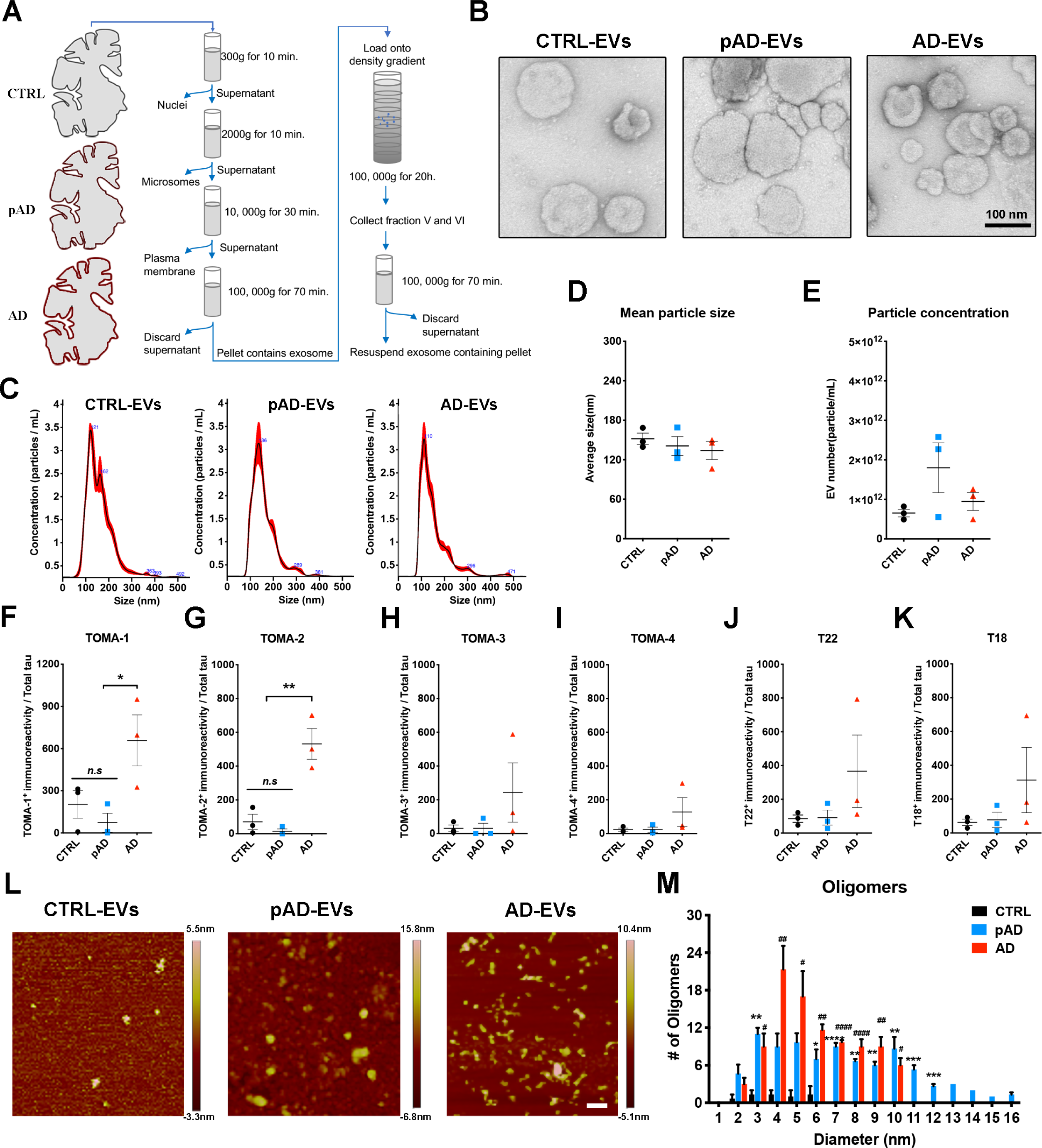
Characterization of EVs by TEM, nanoparticle tracking analysis, tau oligomer dot-blotting and atomic force microscopy. **a.** A schema of EV separation from human frozen brain tissue. **b.** TEM image of human brain-derived EVs. **c-e.** Nanoparticles tracking analysis (NTA) of isolated EVs (C), quantification of EV size (D) and EV density (E). **f-k**. Semi-quantification of tau oligomers in EVs by multiple tau oligomer antibodies. Dot blot images shown in Supplementary Fig. 1. ^**\***^*p* < 0.05, ^**\*\***^*p* < 0.01, as determined by one-way ANOVA (alpha = 0.05) and Turkey’s *post-hoc*. Graphs indicate mean ± s.e.m. Each dot represents individual subject, 3 replicates per subject, 3 subjects per group. **l-m.** Atomic force microscopy (AFM) images showing brain-derived EV-tau oligomers isolated from CTRL, pAD, and AD brains (L), scale bars = 200 nm. Size distribution histogram of EV-tau oligomers (M). ^*^*p* < 0.05, ^**^*p* < 0.01, ^***^*p* < 0.005 and ^****^*p* < 0.0001 for pAD EVs vs. CTRL EVs; ^#^*p* < 0.05, #^#^*p* < 0.01, and ^####^*p* < 0.0001 for AD EVs vs. CTRL EVs as determined by one-way ANOVA (alpha = 0.05) and Turkey’s *post-hoc*. Graphs indicate mean ± s.e.m. n=3 images per sample.

### Increased uptake of AD EVs by primary neurons leading to tau transfer

Numerous mechanisms of EV uptake have been proposed including endocytosis, macropinocytosis, phagocytosis, caveolae-dependent, clathrin-dependent, lipid raft-dependent endocytosis and membrane fusion [68]. Protein-protein interaction between EVs and cell surface molecules on the recipient cells can facilitate the binding of EVs and subsequent endocytosis [50, 68]. For example, interaction between the integrin family on EVs and intercellular adhesion molecules (ICAMs) [49] or extracellular matrix, including fibronectin and laminin, on the recipient cell surface are important for EV binding [58, 67, 68]. Furthermore, heparin sulfate proteoglycan (HSPG) [15] and galectin [7] can mediate EV uptake. We hypothesized that disease conditions alter molecular complex of EV surface and change the efficiency of EV uptake. We tested brain derived-EVs for their uptake by primary cultured murine neurons *in vitro*. After seven days of neuronal differentiation, the cells were incubated for 24 hours with tau containing PKH26-labeled EVs isolated from the brain tissue of AD, pAD and CTRL samples (AD-EV tau, pAD-EV tau, and CTRL-EV tau) and examined for the EV uptake as previously described [74] (Fig. 2a). The neuronal EV uptake was significantly higher in AD EVs compared to CTRL EVs, while the neuronal uptake of pAD EVs was similar to CTRL EVs (Fig. 2b-c). Concomitantly, the transfer efficiency of tau from EVs to neurons normalized by the original tau input was significantly higher in AD EVs compared to CTRL EVs (Fig. 2d). We labeled the supernatant with PKH26 as a negative control at the last ultracentrifugation wash-step of the EV isolation and applied it to neuronal cells. There was no PKH26 positivity found in supernatant-applied neurons (data not shown). EV surface protein repertoires are known to reflect their biological condition and cell-type specificity of parental cells [39, 68]. Our recent proteome of human brain-derived EVs revealed that CTRL EVs expressed more protein of neuronal origin while AD EVs showed more glial dominance. This may reflect the neuroinflammatory condition recently described as the third core AD pathogenesis following Aβ plaques and neurofibrillary tangles [41]. Thus, these results corroborate an idea that a selective AD EV surface molecules may facilitate their uptake by recipient neurons. Finally, to understand if EV-tau has different tau seeding activity dependent on the disease conditions, we employed a FRET sensor-based tau seeding assay as previously described [36]. Astonishingly, the AD EVs showed significantly higher seeding activity compared to pAD and CTR-EVs group (Fig. 2e), suggesting higher potency of AD EVs to induce tau pathology. In summary, the data demonstrate pathogenic functions of AD EVs with efficient transfer of tau and high seeding potency.

**Figure 2.**
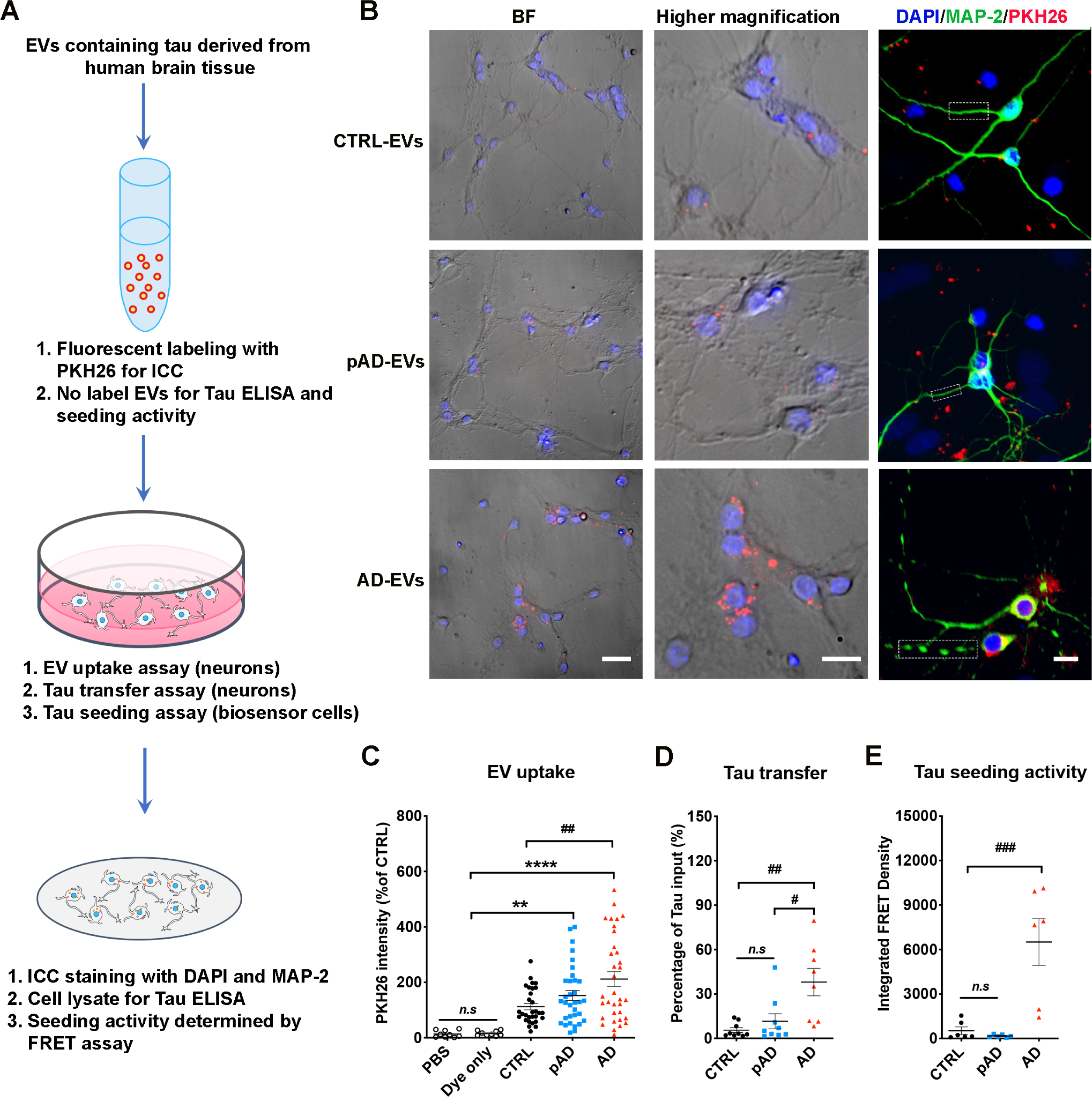
Neuronal uptake, tau transfer efficiency and tau seeding activities of human brain-derived EVs. **a.** A diagram illustrating the primary culture model with primary neurons employed to measure the transfer of EVs containing tau and a biosensor cell assay system for seeding activity. **b.** Cellular uptake of PKH26-labeled EVs (red) by primary culture murine cortical neurons (MAP-2, green; DAPI, blue). Original magnification: 20× (left and middle columns); 40× (right column, taken by Zeiss LSM710 confocal microscopy). Scale bars = 40, 20, 10 µm (left to right). **c.** Quantification of PKH26 fluorescent intensity in MAP-2^+^ neurons. ^**^*p* < 0.0001 and ^****^*p* < 0.0001 compared with PBS or Dye only group; ^##^*p* < 0.01 compared with CTRL-EV group; determined by one-way ANOVA (alpha = 0.05) and Turkey’s *post-hoc*. Each dot represents average data per cell in one image (10-20 cells per image), 30 images per group (for PBS and dye only), 10 images per donor and three donors per group (for CTRL-EV, pAD-EV and AD EVs), total N = 30 per group. **d.** Total human tau ELISA of neuronal cell lysates. ^#^*p* < 0.05 compared with pAD-EV and ^##^*p* < 0.01 compared with CTRL-EV group; *n.s* denotes no significance as determined by one-way ANOVA (alpha = 0.05) and Turkey’s *post-hoc*. Three donors per group, three independent experiments. Graphs indicate mean ± s.e.m. **e.** EVs were tested in the Tau-FRET assay for tau seeding activity. Results are plotted as integrated FRET Density values for each sample. ^###^*p* < 0.001 compared with CTRL-EV and pAD-EV group; as determined by one-way ANOVA (alpha = 0.05) and Turkey’s *post-hoc*. Three donors per group, and each dot represents one well. Graphs indicate mean ± s.e.m. **b-e**: Three donors per group, and the data is representative of three independent experiments.

### Inoculation of AD EVs propagate tau pathology in aged mice

Since we observed efficient EV uptake and transfer of EV tau into primary mouse cortical neurons and significant tau seeding activities in AD EVs, we further tested whether brain derived EVs can initiate tauopathy in 2 months-old B6 mice after an intrahippocampal injection. Brain derived EVs containing tau isolated from the brain tissue of AD, pAD, and CTRL cases were unilaterally injected in the OML of the DG (Fig. 3a). The amount of injected tau (300 pg/μl, 1ul injection) was much lower than the one used for the previous tau propagation studies (1-8 μg) [33, 54]. Its concentration was in a range of the extracellular tau concentration in mouse interstitial fluid of the central nervous system [73]. Immunofluorescence against phosphorylated tau (p-tau) antibody AT8 (pSer202/pSer205) detected a considerable, yet not an abundant, amount of AT8^+^ cells in the hippocampal region of AD and pAD EVs injected female mice (Supplemental Fig. S2, left) but not in male mice (data not shown). A previous study reported a more enhanced tau propagation induced by fibril tau injection with aged B6 mice in comparison to young mice [33]. Therefore, we decided to use aged female mice as recipients to determine if tau pathology induced by brain-derived EVs reflects the donor’s disease conditions. Brain-derived EVs were isolated from 2 donors of each AD, pAD, and CTRL cases and from *Mapt* knockout (Tau KO) mice as the control. Each EV sample (containing 300 pg tau/injectate for human brain derived EVs), or saline as an injection control, were unilaterally injected into the OML of the DG of ∼18-months-old B6 female mice (Fig. 3a). The spread of tau pathology was evaluated by immunofluorescence against AT8 in the hippocampal region at 4.5 months post injection (Fig. 3b, Supplementary Fig. S2, right, and S3a). Interestingly, abundant perikaryal AT8^+^ inclusions were detected in both ipsilateral and contralateral sides of the hippocampal region including the Cornus Ammonis 1 (CA1), CA3, dentate granule cells, subgranular zone, and hilus in the AD and pAD EVs groups, suggesting tau transfer between anatomically connected pathways (Fig. 3b). Semiquantitative brain-wide mapping of tau pathologies revealed that AT8^+^ pathogenic tau was accumulated throughout the hippocampus, predominantly distributed in the caudal hippocampal hilus, in the mouse brains injected with AD or pAD EVs, while CTRL EVs injected mouse brains showed very little AT8 positivity (Fig. 3c). Notably, the percentage of the area occupied by AT8^+^ cells in the hippocampal region was significantly higher in AD EVs as compared to CTRL EVs, Saline or Tau KO EVs groups (Fig. 3d). There was no significant difference between pAD and CTRL EVs injected groups and no AT8^+^ staining was observed in saline group (Fig. 3d, Supplementary Fig. S3a). All AT8^+^ neurons were negative for human tau as determined by immunofluorescent staining against human tau-specific monoclonal HT7 (data not shown), indicating that endogenous mouse tau was recruited and aggregated by the inoculation of human brain derived EV tau. A growing body of evidence suggests that misfolded tau tends to be truncated and frequently consists of different conformers or structural polymorphisms, deciphering the stages and disease of tauopathy [25, 26, 29, 64, 77]. Therefore, we performed neuropathological analysis of tau by immunohistochemistry using conformation-specific (Alz50 and MC1) and p-tau epitope-specific monoclonal (CP13: pSer202 tau, PS422: pS422 tau, and PHF1:pSer396 and pSer404). All 5 antibodies detected misfolded or phosphorylated tau mainly in the hilus of hippocampal region with AD and pAD EV groups (Supplemental Fig. S3b-f).

**Figure 3.**
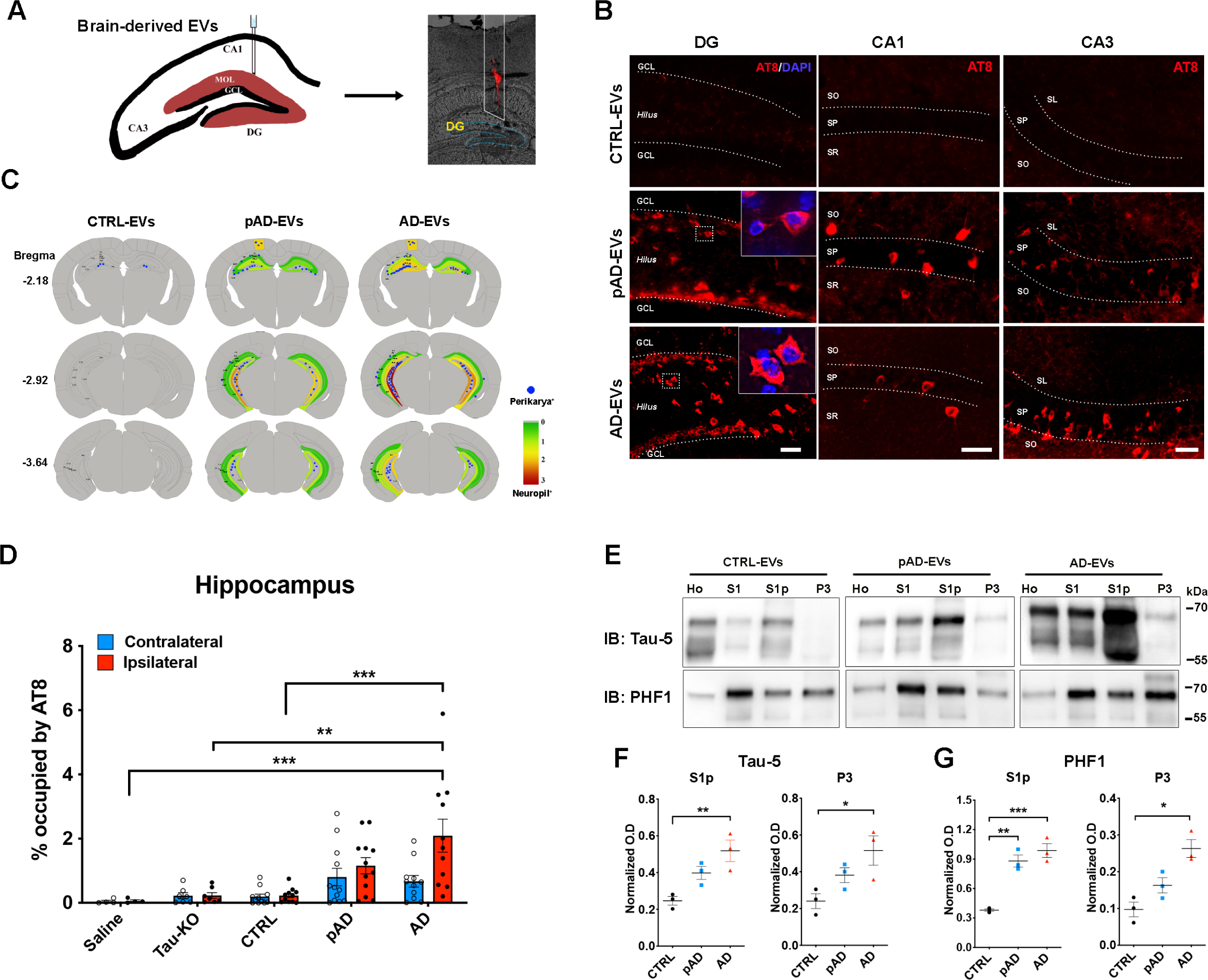
AD-EV but not CTRL-EV injection causes progressive tauopathy in aged B6 mouse brains. **a.** A schema illustrating 300 pg of tau containing EVs from human brain unilaterally injected to the hippocampus of B6 mice at 18-19 months of age. DiI (red) indicated the injection site of outer molecular layer of hippocampus. **b.** Representative image of AT8 staining (red) 4.5 months after intrahippocampal injection of AD EV and pAD EV into aged B6 mouse brain. Original magnification: 20×, Scale bar = 50 μm. **c.** Semiquantitative analysis of AD-like tau pathologies based on AT8 immunostaining of brains from CTRL-EV, pAD-EV and AD-EV-injected mice at 4.5 months post injection. Blue dots represent AT8^+^ perikaryal inclusions. AT8^+^ density from green (0, low) to red (3, high). **d.** Quantification of AT8+ occupied area in the contralateral (blue) and ipsilateral (red) in entire hippocampal regions of recipient mice. ^*^*p* < 0.05 and ^**^*p* < 0.01 compared with CTRL-EV group determined by one-way ANOVA (alpha = 0.05) and Turkey’s *post-hoc*. Total mice in each group for the quantification are 4, 6, 12, 12, 11 for saline, Tau-KO, CTRL, pAD and AD. Two donors for EVs per group for CTRL, pAD and AD (n = 5-6 mice per donor). Bregma −1.34 to −3.64, 4 sections per mouse were analyzed. Each dot represents mean value from one animal. Graphs indicate mean ± s.e.m. **e.** Immunoblotting of biochemically fractionated brain tissue samples for homogenate (Ho), TBS supernatant (S1), tau oligomer enriched (S1p) and tau fibril enriched fractions (P3) by Tau-5 (total tau) and PHF1 (pSer396/pSer404 tau) (top panels) and their quantification (bottom panels). Equal proportions of Ho, S1, Sp1 and P3 fractions were analyzed (n = 3 mice / group). Optical density (OD) was normalized to that for the homogenate fraction from each corresponding mouse. ^*^*p* < 0.05 and ^**^*p* < 0.01 compared with CTRL group as determined by one-way ANOVA (alpha = 0.05) and Turkey’s *post-hoc*. Graphs indicate mean ± s.e.m.

We next examined whether EV-tau could induce templated misfolding of original tau aggregates in endogenous tau of the recipient mice. Aggregated tau was extracted from the recipient mouse brains via sarkosyl solubilization and sequential centrifugation, and immunoblotted using Tau-5 and PHF1 monoclonal antibodies as previously described (Fig. 3e) [1, 40]. We observed a significant increase in oligomeric tau in the fraction S1p of both AD and pAD EV injected mouse hippocampi as compared to the CTRL EV group, determined by both Tau-5 (total tau) and PHF1 immunoblotting (Fig. 3f-g). The amount of sarkosyl-insoluble tau in the fraction P3 was also significantly elevated in AD EV injected mouse hippocampi when compared to CTRL EV group (Fig. 3f-g). These data indicate that AD EV inoculation induced accumulation of oligomeric and fibrillar tau, while pAD EV inoculation induced accumulation of oligomeric tau. Taken together, these data show the efficient induction of tau propagation in the hippocampus of the aged B6 female mouse brain after the injection of the AD EVs containing physiological concentration of tau. Conformational changes of tau in the recipient mice appear to reflect the original tau confirmation of AD EVs and pAD EVs, which were also reported with mice injected with AD brain-derived tau fibrils [33].

### Inoculation of AD EVs show more tau propagation as compared to the inoculation of an equal amount of tau oligomer or fibril-enriched fractions from the same AD brain tissue

To determine how propagation of tau pathology may differ between the injection of EV-associated or free form tau, we compared EV tau with oligomer and fibrillar tau derived from the same donor for tau pathology development. Fibril or oligomeric tau were isolated from the same AD EV donor as S1p and P3 fractions according to the previous publications [1, 33, 40]. The p-tau immunoreactivity and structure of the isolated tau aggregates were examined by the western blot using PHF1 antibody and AFM (Fig. 4a-b). AFM images showed mostly small oligomer like globular particles (6-8nm in height) in EV and tau oligomer preparation and large globular structures (30-70nm in height) in sonicated tau fibril preparations (Fig. 4a), which is consistent with the description of the fibril structure as previously reported [30]. We observed mainly monomeric PHF1^+^ band in p-tau in EV and tau oligomer enriched samples, and trimeric PHF1^+^ band in fibril enriched sample (Fig. 4b), validating their oligomeric and fibrillar conformation. We injected each sample of AD EV, oligomers, and fibrils containing an equivalent amount of tau (300pg / 1μL injectate) into the OML of the DG of 18-month-old B6 female mice. At 4.5 months after the injection, mice were euthanized and tested for tau pathology by immunofluorescence against AT8. We observed strong AT8 positivity in the injection site with all groups, suggesting successful intrahippocampal injections. In addition, AT8^+^ signal was also seen as perikaryal inclusions or neuropil staining in the cortex along the needle tract (Fig. 4c, top panels, Supplementary Fig. S4, left), whereas only neuropil accumulation of tau with oligomer or fibril tau injected mice, which is in agreement with the previous study [33] (Fig. 4c, top panels, Supplementary Fig. S4, middle and right). Moreover, compared to AT8^+^ tau pathology observed in the entire hippocampal region with AD EV injected mice as described previously, fibril or oligomer tau injected mice did not show any AT8^+^ perikaryal inclusions in the entire hippocampus (Fig. 4c, bottom panels, d). Consistent with the previous reports [33, 44], injecting 2 μg of oligomer or fibril tau from AD brain tissues in the aged B6 mice induced robust tau pathology in the hippocampal region, thus providing the fidelity of our oligomer or fibril tau isolation methods (Supplementary Fig. S5a-c). These findings recapitulated our previous study showing that inoculation of microglia-derived EVs containing 5ng of aggregated tau, but not inoculation of the equal amount of free tau aggregates, was able to induce tau propagation in the DG of B6 mice [4]. Previous studies reported that inoculation of 1-8 μg of fibril tau from AD patients into wildtype (B6 and B6/C3H F1) mouse brains could induce tau propagation as early as 3 months post injection [33, 54]. Potency of propagation may be varied between the donors and the type of tauopathies [54], therefore it is difficult to compare the results between these studies. To the best of our knowledge, this is the first report of increased tau propagation potency in EV-tau as compared to vesicle free tau isolated from the same human AD brain tissue. A previous study reported that immunodepletion of tau from the AD brain derived tau fibril diminished tau aggregation activity *in vitro* or propagation *in vivo* [33]. Moreover, addition of remaining components after the immunodepletion of AD-tau into fibril tau did not alter the outcome of abovementioned experiments, suggesting that tau was the essential component to initiate tau propagation but not tau associated molecules [33]. Our results also indicated that EVs without tau do not initiate tau propagation *in vivo* as we barely observed tau propagation by injecting Tau KO EVs. We, however, confirmed that EVs certainly enhanced propagation potency of tau. The discrepancy between these experiments may be due to the potential removal of EVs associated with extracellular tau when immunodepletion of tau was performed.

**Figure 4.**
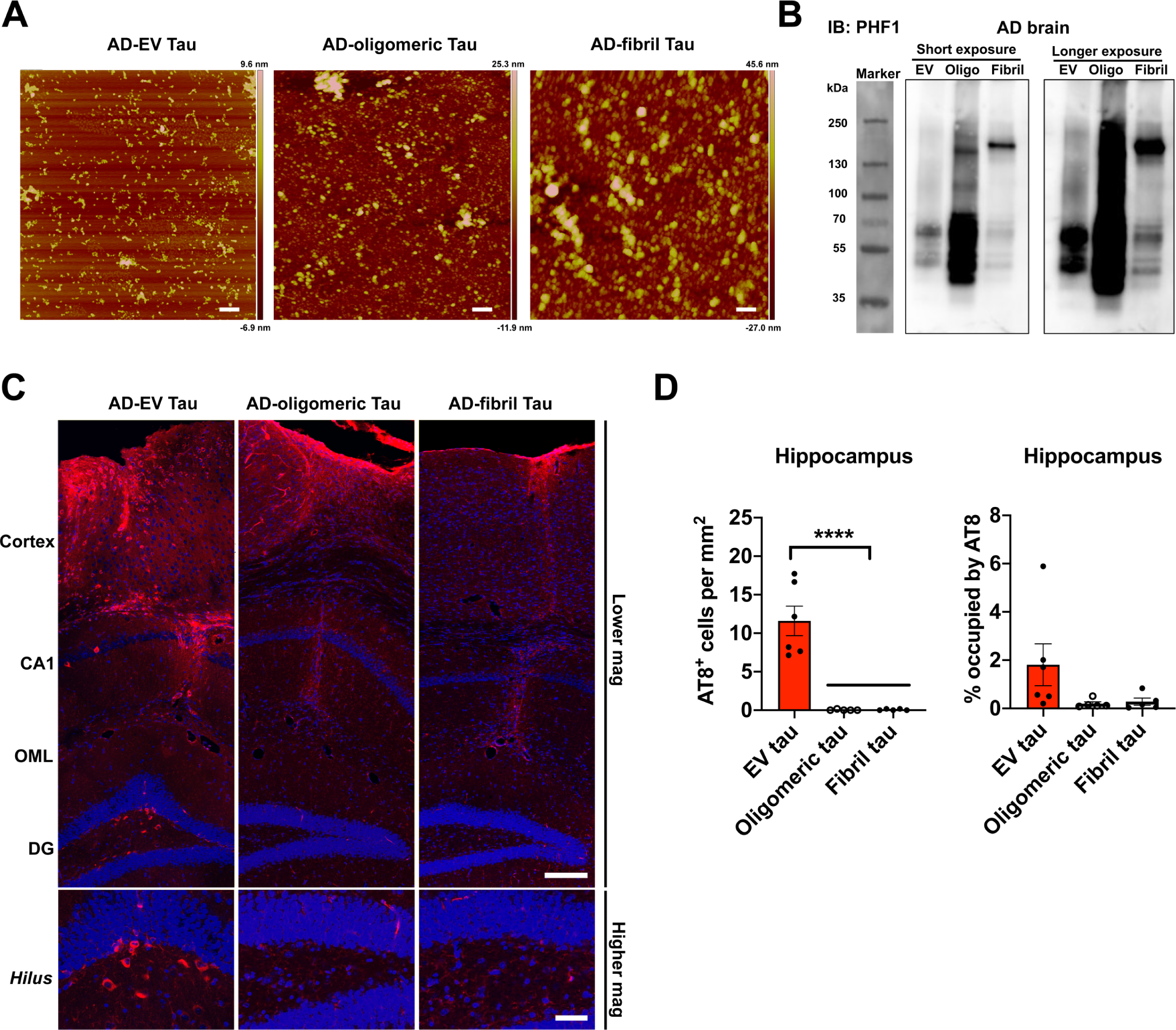
EV-tau but not oligomeric or fibril tau enriched samples derived from the same AD brain induced tau propagation in mouse brain. **a.** AFM images of EVs and tau aggregates isolated from the same AD brain tissues. Scale bars = 200 nm **b.** Representative images of PHF1 immunoblotting of isolated EVs, tau oligos and tau fibrils by PHF1 antibodies. **c.** Representative images of AT8 immunostained recipient mice after unilateral injection of AD EVs (left), tau oligomer-enriched fraction (middle) and tau fibril-enriched fraction (right) in cortical region (top panels) and dentate gyrus (bottom panels). Scale bars = 200 µm (top), 50 µm(bottom). **d.** Quantification of AT8^+^ neurons in the hippocampus of recipient mice. ^****^*p* < 0.0001 compared between EV-tau injected group and oligomeric or fibril tau group, as determined by one-way ANOVA (alpha = 0.05) and Turkey’s post-hoc. EV-tau, oligomeric and fibril tau group: n = 5-6 mice per group for quantification. Bregma −1.34 to −3.64, 4 sections per mouse were analyzed. Each dot represents mean value per animal. Graphs indicate mean ± s.e.m.

### Preferential EV-mediated tau propagation to GABAergic inhibitory neurons

Recent work indicates that specific type of organs or cells, where EVs are transferred, could be determined by the enriched proteins on the EV surface [68]. For example, previous studies found that specific EV proteins, such as integrins or tetraspanins, play critical roles for the deliveries of cancer-derived EVs to specific organs or cell types [37, 55]. Given the fact that some EV surface proteins are specifically expressed on AD EVs [51], we speculated that the evaluation of EV-mediated transfer of tau to aged mouse brains would uncover cell type-specific tau transfer mechanisms. To determine which neuronal cell type preferentially accumulates tau, we performed double immunostaining using the markers for p-tau (AT8) and GABAergic interneurons (GAD67 and parvalbumin, PV) or excitatory neurons (Neurogranin, NG, and glutamate receptor 2/3, GluR2/3, mossy cell marker) [69]. Surprisingly, most of AT8^+^ cells were GAD67^+^ interneurons in the CA1, CA3, and DG region in AD EV and pAD EV injected mice (Fig. 5a-b). Moreover, a subset of PV^+^ neurons were also co-localized with AT8 (Supplementary Fig. S6a). We found that the ratio of GAD67^+^AT8^+^ cells over total GAD67^+^ cells were significantly higher in the DG and CA3 region in AD EV and pAD EV, and in the CA1 in AD EV compared to CTRL EV injected mice (Fig. 5c-e), although there was no significant reduction in the total number of GAD67^+^ neurons in those regions. No difference was observed in any of the regions between Tau KO EV and CTRL EV groups. In contrast, no NG^+^ excitatory neurons were AT8^+^ in the DG of hippocampus (Supplementary Fig. S6b). We, however, observed that some of AT8^+^ cells were GluR2/3^+^ mossy cells in the hilus region (Fig. 5f). Quantification of AT8^+^ cells in the hippocampal region in AD EV injected mice revealed that 64% and 23% of the AT8^+^ cells were GAD67^+^ inhibitory neurons and GluR2/3^+^ excitatory mossy cells, respectively (Fig. 5g). Multiple lines of evidence have supported the notion that GABAergic interneuron dysfunction could be one of critical components in the early pathogenesis of AD. The decreased levels of GABA transmitter have been reported in the CSF of AD patients or elderly without cognitive impairment [6, 78] and in their post-mortem tissues especially in the temporal cortex, followed by the hippocampus, frontal cortex, and thalamus of AD patients [31]. AD patients showed loss of specific somatostatin^+^ interneurons in the hippocampus and cortex [13, 19]. Moreover, 7-21% of sporadic AD patients show at least one episode of seizure during the illness [56], and administration of anti-epileptic drug, levetiracetam, was effective to improve cognitive function in the elderly for those with normal memory, MCI, and AD patients [62, 70]. Together, our data indicate that EVs may play a critical role in tau propagation to GABAergic neurons, and suggest that EVs can be an attractive therapeutic target for the early intervention of AD.

**Figure 5:**
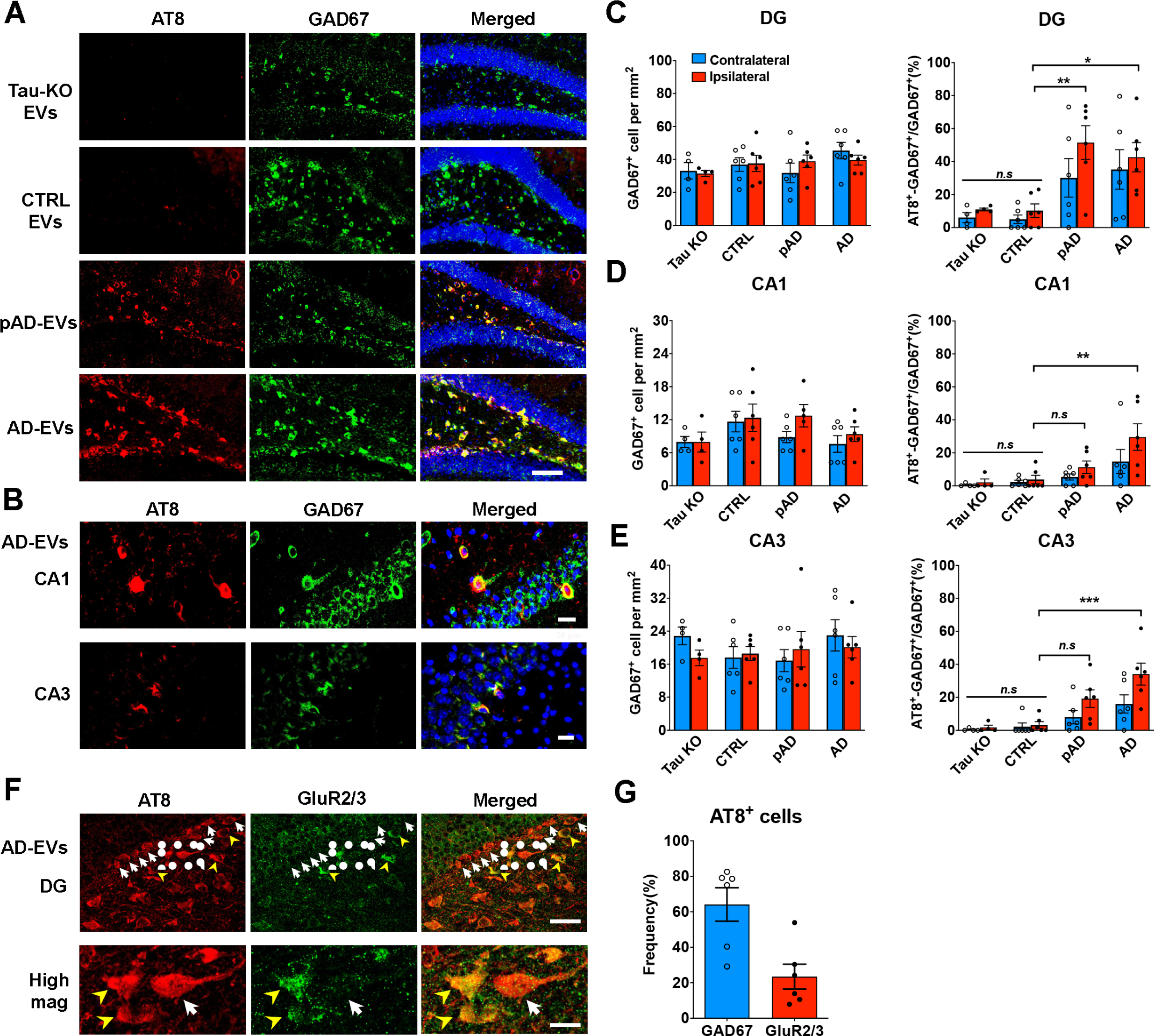
Specific pathological tau staining with AT8 antibody in GABAergic interneurons in the hippocampus of B6 mice. **a.** AT8 (red) and GAD67 (green) immunostaining in the ipsilateral dental gyrus of hippocampal region from Tau KO EV, CTRL EV, pAD EV and AD EV injected mice at 4.5 months post injection. Scale bars = 100 µm. **b.** AT8 (red) and GAD67 (green) immunostaining in the ipsilateral CA1 and CA3 of hippocampal region from AD EV injected mice. Scale bars = 20 µm(top), 25 µm (bottom). **c-e.** Quantification of GAD67^+^ cells in DG (c), CA1 (d) and CA3 of hippocampus (e). The percentage of AT8^+^ GAD67^+^ cells in all GAD67^+^ cells are shown in the right column (c-e). Ipsilateral side (red column) contralateral side (blue column)^. *^*p* < 0.05, ^**^*p* < 0.01 and ^***^*p* < 0.001 compared with CTRL group, as determined by one-way ANOVA (alpha = 0.05) and Turkey’s *post-hoc*. n = 5-6 mice per group for quantification. At least two sections were imaged per animal. Each dot represents mean value per animal. Graphs indicate mean ± s.e.m. **f-g.** Immunostaining of GluR2/3^+^ mossy cells (f) and AT8 in the ipsilateral dentate gyrus of hippocampal region from AD-EV injected mice; and quantification of the ratio of GAD67^+^ AT8^+^ cells / total AT8^+^cells (blue) and GluR2/3^+^ AT8^+^ cells / AT8^+^ cells (red) (g). n = 6 mice per group for quantification. At least two sections were imaged per animal. Each dot represents mean value per animal. Graphs indicate mean ± s.e.m. Scale bars = 20 µm(top), 10 µm(bottom).

### AD EV and pAD EV inoculation reduced GABAergic neuronal activity and input to CA1 pyramidal cells

To determine if EV-mediated tau propagation may disrupt GABAergic neuronal functions, we examined the neuronal activity of GAD67^+^ GABAergic neurons by immunofluorescence against c-fos. There was a significant reduction in c-fos^+^/ GAD67^+^ cells in the CA1 in AD EV as compared to Tau KO EV injected mice (Fig. 6a-b). However, there was no significant difference in c-fos^+^/ GAD67^+^ cells in the DG between any groups (Fig. 6c-d), suggesting decreased neuronal activity in GABAergic neurons specifically in the CA1 region by EV-mediated tau propagation. We further assessed the synaptic input of GABAergic neurons to CA1 pyramidal cells by examining the number of immunostained GAD67^+^ puncta surrounding CA1 pyramidal neuronal cell soma. The images were captured by confocal microscope and the number of the puncta was analyzed by Imaris software (Fig. 6e). There was a significant reduction in the number of puncta in the CA1 pyramidal layer of pAD EV and decreased tendency with AD EV as compared to Tau KO EV group (Fig. 6f). There was no difference between the groups in the cell numbers of CA1 pyramidal neurons (Fig. 6g). CA1 pyramidal neurons receive abundant inhibitory inputs from GABAergic neurons [12], therefore, our results suggest possible dysregulated function in CA1 pyramidal neurons via disrupted GABAergic neuronal function after EV-mediated tau propagation.

**Figure 6.**
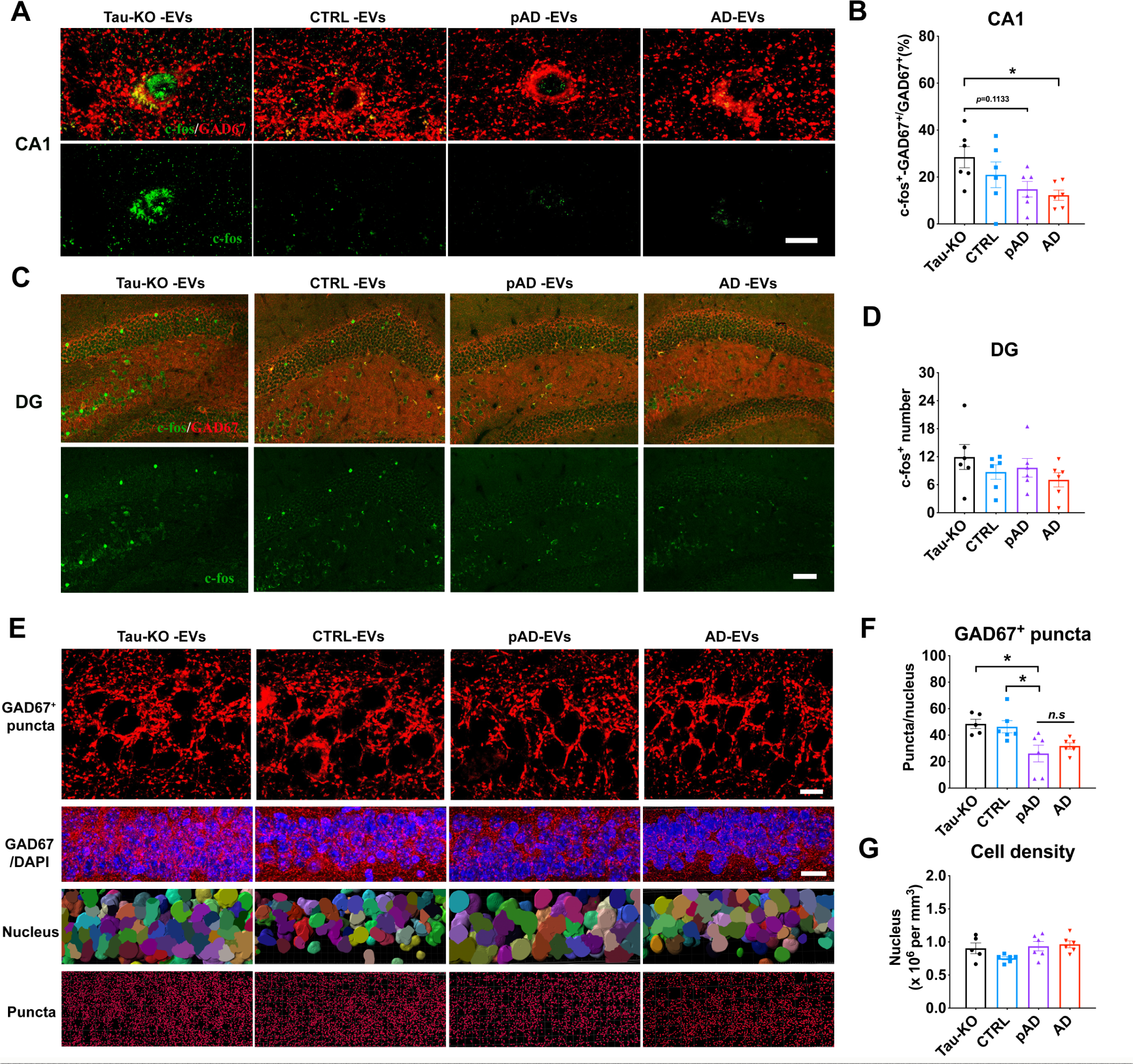
Reduction in c-fos expression in GAD67^+^ GABAergic neurons and GAD67^+^ puncta around CA1 pyramidal cells in AD EV and pAD EV injected aged B6 mice. **a-b.** GAD67 (red) and c-fos (green) co-staining images (a) and quantification of the percentage of c-fos^+^ GAD67^+^ cells in all GAD67^+^ cells (b) in CA1 region. Scale bar=10 μm. **c-d.** GAD67 (red) and c-fos (green) co-staining images (c) and quantification of the percentage of c-fos^+^ GAD67^+^ cells in all GAD67^+^ cells (d) in DG region. Scale bar=50 μm. ^*^*p* < 0.05 AD-EVs compared with Tau-KO EV group, as determined by one-way ANOVA (alpha = 0.05) and Turkey’s *post-hoc*. n = 6 mice per group for quantification. At least two sections were imaged per animal. Each dot represents mean value per animal. Graphs indicate mean ± s.e.m. **e.** High-magnification images in top panels compared GAD67 expression (red) in CA1 pyramidal cells of hippocampus all four injected Tau-KO-, CTRL-, pAD- or AD EV groups. Scale bar=10 μm. Second panel shows lower-magnification images of GAD67 expression and DAPI staining. Scale bar=20 μm. Third panel shows cells counted by Imaris software based on DAPI staining. Fourth panel shows GAD67+ puncta analysis by Imaris. Scale bar: 10 μm. **f-g.** Quantification of GAD67^+^ puncta (f) and total cell number in CA1 of hippocampus (g). ^*^*p* < 0.05 and ^**^*p* < 0.01 pAD-EV compared with Tau-KO and CTRL-EV group, as determined by one-way ANOVA (alpha = 0.05) and Dunnett’s *post-hoc*. n = 5-6 mice per group for quantification. At least two sections were imaged per animal. Each dot represents mean value per animal. Graphs indicate mean ± s.e.m.

### EV-induced alterations in intrinsic membrane properties and spontaneous inhibitory synaptic currents in CA1 pyramidal neurons

To evaluate the functional effect of tau propagation in human brain-derived EV-inoculated mouse brains, we performed whole-cell voltage/current–clamp recordings of CA1 pyramidal cells using 300 µm-thickness acute tissue slices of mouse hippocampi from Tau KO EV, pAD EV, and AD EV groups (Fig. 7a-b). An F-I curve-generating protocol ranging from −100 pA to +120 pA square pulse current steps (increments of +20 pA) or −220 pA to +330 pA current steps (increment of +50 pA) were applied. The number of action potentials (APs) evoked by depolarizing current steps was significantly lower in pAD EV and AD EV groups compared to Tau KO EV groups as determined by repeated measurement ANOVA (Fig. 7c, e, Supplementary Table S4-5) and for pAD EV group compared to Tau KO EV group at +100 pA (p=0.0434) and +130 pA (p=0.0445) (Fig. 7d, f). This result is consistent with the study on another tau transgenic mouse model (aged rTg4510 mice expressing P301L tau), which show reduction in firing in hippocampal CA1 neurons [34]. The AD EV group also showed significant reduction of mean AP amplitude as compared to Tau KO EV group (Fig. 6g). Evaluation of the properties of spontaneous inhibitory and excitatory postsynaptic potentials (sIPSCs and sEPSCs) (Fig. 7h and Supplementary Table S6-8) revealed a significant reduction in the mean amplitude of sIPSCs in pAD EV group and E-I ratio of amplitude as compared to Tau KO group (Fig. 7h-i). There was no difference in sEPSC properties among the 3 groups. Taken together, these data demonstrate reduction in action potential firing rates of CA1 pyramidal neurons in the pAD EV group, reduction of AP amplitude in the AD EV group, and reduction in sIPSC amplitude in the pAD EV group, which is also reflected in the reduction in the E-I ratio of sIPSC amplitude. Thus, pathogenic tau accumulation may compromise both intrinsic excitability (evoked action potential firing rates) and inhibitory synaptic responses of CA1 pyramidal cells.

**Figure 7.**
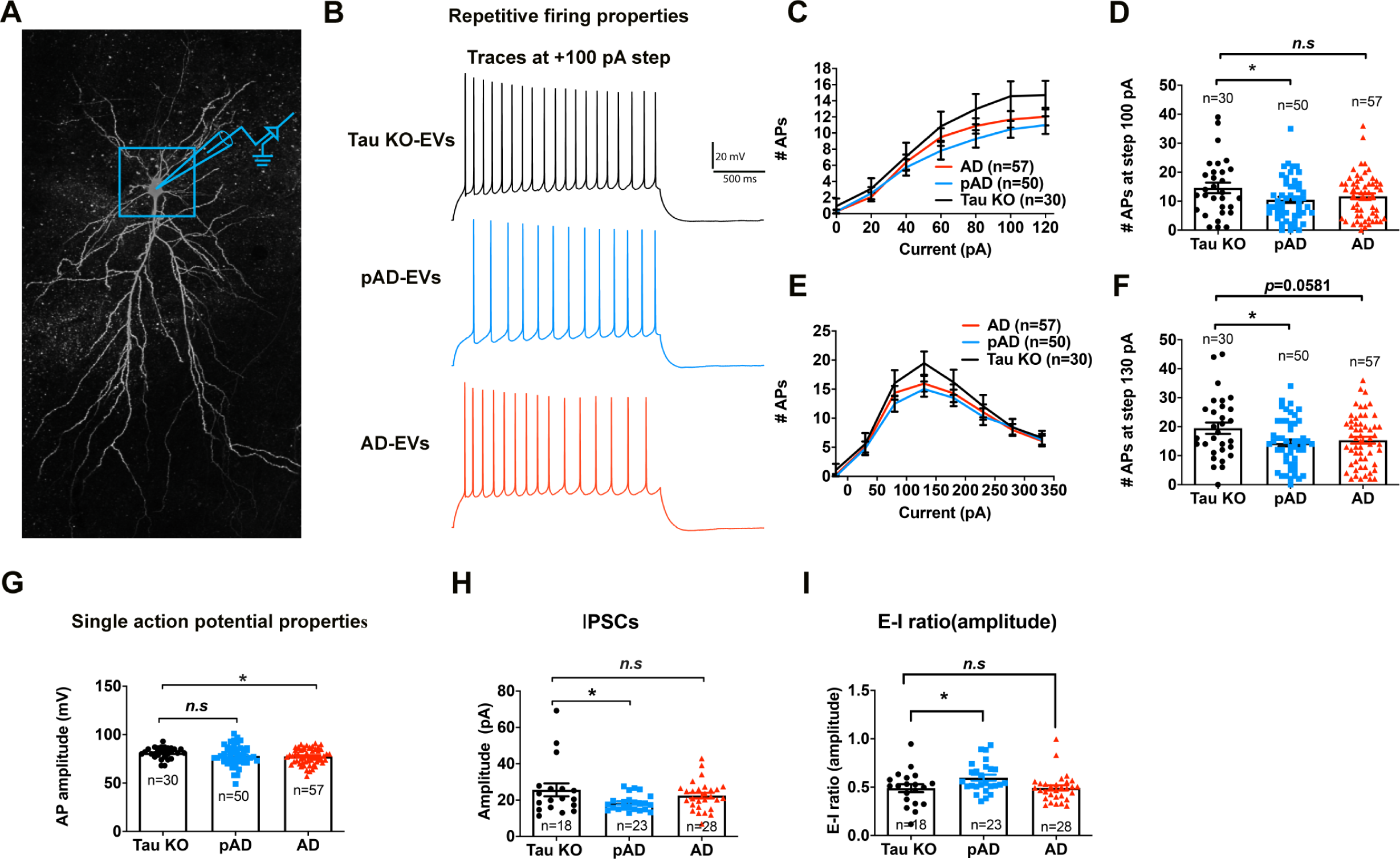
whole-cell current clamp recording of CA1 pyramidal neurons. **a**. Confocal z stack montage (63× magnification) image of biocytin-filled mouse CA1 pyramidal neurons after recording. **b-g**: Action potential (AP)-firing recorded in whole-cell current clamp mode; **b**: Representative traces for Tau KO (black color), pAD (blue color), and AD-EV (red color) for 100 pA steps at 2 s long High Rn protocol. **c**. Quantification of repetitive firing at High-Rn step current injection protocol. ^**^p<0.01 vs. Tau KO-EV group as determined by RM-ANOVA; **d**: pAD-EV significantly reduce the firing at 100 pA; **e**: Quantification of repetitive firing at Low-Rn step current injection protocol. ^*^p<0.05 vs. Tau KO-EV group as determined by RM-ANOVA; **f**: pAD-EV significantly reduced the firing rate at and + 130 pA of step current; **g**. AD-EV significantly reduced AP amplitude. **c-g**: n = 30, 50, and 57 cells for Tau KO, pAD and AD-injected mice, 5-7 mice per group. Each dot represents one recorded cell. Graphs indicate mean ± s.e.m. **h-i**: Quantification of GABAergic spontaneous inhibitory postsynaptic currents (sIPSCs) recorded in whole-cell voltage clamp mode from neuronal network. pAD showed significant decrease in sIPSC amplitude (h) and E-I amplitude ratio (i). ^*^*p* < 0.05 compared with CTRL group, as determined by one-way ANOVA (alpha = 0.05) and Dunnett’s *post-hoc*. H-I: n = 18, 23, and 28 cells for Tau KO, pAD and AD-injected mice, 5-7 mice per group. Each dot represents one recorded cell. Graphs indicate mean ± s.e.m. See also Supplementary Tables S4-S7.

## Discussion

The current study demonstrated that AD EVs efficiently initiated tau propagation in aged B6 mice. This finding was validated by the *in vitro* evidence of the highly transmissible nature of AD EVs with their higher uptake by cortical neurons and increased seeding activity compared to CTRL EVs. Tau pathology was predominantly found in GABAergic neurons and to a lesser extent in mossy cells in the DG. Whole-cell patch clamp recording of CA1 pyramidal cells of recipient mice showed reduced intrinsic excitability and lower mean sIPSC amplitude indicative of intrinsic dysfunction of CA1 pyramidal cells and reduced input from interneurons. This was accompanied with reduced inhibitory synaptic markers and c-fos immunoreactivity in GABAergic neurons in the CA1 region. The preferential EV mediated tau propagation into GABAergic neurons and their reduced function posits the potential underlying mechanism in interneuron dysfunction in AD.

Recent advances in EV research have opened new avenues to investigate the diagnostic and pathogenic roles of EVs on neurodegenerative diseases [21, 22, 75]. Accumulating evidence now suggests that EVs carry pathogenic proteins, and EV-associated proteins or miRNAs predict disease progressions in AD [14, 72], chronic traumatic encephalopathy [66], Parkinson disease, prion disease, amyotrophic lateral sclerosis, traumatic brain injury, multiple sclerosis, and Huntington disease [22, 75]. Furthermore, overexpression of the second most AD-associated GWAS gene, Bridging integrator-1 (BIN1), enhanced release of tau via EVs *in vitro* and exacerbated tau pathology in PS19 mice *in vivo* [47]. Contribution of EVs to tau pathology development in AD patients has been questioned, however, due to the scarcity of tau in the EV fractions of biofluids. We have demonstrated here that EVs containing only 300 pg of tau successfully induced templated misfolding in endogenous tau and subsequently transferred tau pathology through the entire hippocampus in aged B6 mice, indicating that EVs are indeed vehicles to transfer pathological tau. AD EVs show higher transmissibility of tau via increased uptake by recipient neurons. Our proteome analysis of AD brain-derived EVs suggests enrichment of glia-derived EVs rather than neuron-derived EVs [51]. Interestingly, recent analysis of single cell RNAseq of human AD brains showed that CD81, an established tetraspanin exosome marker, is highly expressed in the microglia module [46] together with ApoE, the most prominent AD GWAS gene [43]. Notably, APOE is a representative disease-associated / neurodegenerative microglia (DAM/MGnD) genes [42], suggesting active EV and APOE synthesis in DAM/MGnD in AD brains. CD81 and CD82 are known to regulate the integrin cluster distribution on plasma membranes to facilitate dendritic cell adhesions [59] and recruit integrins to endosomal pathway [35] respectively. In addition, our recent study demonstrates that IL-1β-stimulated astrocytes secrete EVs enriched in the integrin family with higher neuronal uptake efficiency, which was inhibited by an integrin-blocking peptide [74]. Thus, EV uptake in AD brains could be enhanced by differentially expressed EV surface proteins due to altered cargo sorting or the origin of the cell type in neuroinflammatory conditions.

Dysfunction of interneurons has been extensively reported in tauopathy animal models [45, 65]. JNPL3 transgenic mice harboring *MAPT* P301L mutation show loss of hippocampal interneurons, PHF1^+^ p-tau and MC1^+^ misfolded tau in interneurons, and rescue of enhanced later-phase long-term potentiation by administration of GABA_A_ receptor agonist [45]. VLW mice overexpressing human *MAPT* with 3 mutations (G272V, P301L, and R406W) show p-tau accumulation in hippocampal PV^+^ GABAergic neurons and mossy cells in DG as early as 2 months of age [65]. Reduction of GABAergic septohippocampal innervation of PV^+^ interneurons in VLW mice suggests tau accumulation may be responsible for GABAergic neuronal loss [65]. We found that EV-mediated tau propagation is explicitly in GABAergic neurons, including PV neurons followed by mossy cells, and GABAergic dysfunction was determined by both electrophysiological recording and c-fos activity, indicating the susceptibility of those neurons to tau toxicity. PV neurons are surrounded by the specific extracellular matrix (ECM), called perineuronal nets, comprised of integrin-binding versican and heparin sulfate proteoglycan (HSPG) [20]. Since EV uptake is dependent on HSPG [15], EV surface proteins such as integrins, which are known to interact with HSPG, may play a potential role on their uptake by GABAergic neurons.

In summary, we have revealed the highly transmissible and potent seeding activity of AD EVs with selective susceptibility of GABAergic neurons. Our study created a foundation to elucidate a novel EV-mediated tau spread mechanism, which may be relevant to interneuron dysfunction in AD.

## Materials and Methods

### Animals

Aged C57BL/6 (18-19 months old), Tau KO (B6.129×1-*Mapt*^*tm1Hnd*^/J, # 007251) and pregnant CD-1 mice were purchased from National Institute of Aging (NIA), Jackson laboratory and Charles River Laboratory, respectively. B6 mice were used for intracerebral inoculation of human brain-derived materials. Adult Tau KO mice were used for isolation of brain-derived EVs. E16 CD-1 mice were used for primary culture of cortical neurons. All animal procedures followed the guidelines of the National Institutes of Health Guide for the Care and Use of Laboratory Animals, and were approved by the Boston University Institutional Animal Care and Use Committee (IACUC).

### Isolation of EVs from AD brains

Human and mouse brain-derived EVs were isolated according to our recently published methods [53]. Briefly, fresh frozen human frontal cortex gray matter was sliced with a razor blade on ice while frozen to generate 1–2 cm long, 2–3 mm wide sections. The cut sections are dissociated while partially frozen in 300 μL of 20 units papain (# LK003178, Worthington Biochemical Corporation) in 15 mL Hibernate-E media (Thermo Fisher Scientific) at 37°C for 15 min, and protease and phosphatase inhibitors (# PI78443, Thermo Fisher Scientific) were added. The tissue sample was centrifuged at 300 × *g* for 10 min at 4°C. The pellet was used as the brain homogenate control. The supernatant was centrifuged at 2000 × *g* for 10 min at 4°C. The supernatant was centrifuged at 10,000 × *g* for 10 min at 4°C. The supernatant was transferred through a 0.22-μm filter and ultracentrifuged at 100,000 × *g* for 70 minutes at 4°C using Beckman SW41Ti. The pellet was resuspended in 2 mL of 0.475M of sucrose in double-filtered PBS with 0.22-μm filter (dfPBS) and overlaid on 5 sucrose cushions (2 mL each of 2.0M, 1.5M, 1M, 0.825M, 0.65M in dfPBS) and ultracentrifuged at 100,000 × *g* for 20 h. The samples were fractionated in 1-mL step, and fractions V and VI are collected as EV-enriched fraction. Each fraction was ultracentrifuged at 100,000 × *g* for 70 minutes at 4°C to pellet EVs, which were resuspended in 30 μL dfPBS as a final volume/fraction.

### Nanoparticle Track Analysis (NTA)

The number of EVs in the enriched fraction was analyzed as previously described [52, 53]. Briefly, all samples were diluted in dfPBS for at least 1:1000 or more to get particles within the target reading range for the Nanosight 300 machine (Malvern Panalytical Inc), which is 10-100 particles per frame. Using a syringe pump infusion system (Harvard Laboratories/Malvern), five 60-second videos were taken for each sample at 21°C constant. Analysis of particle counts was carried out in the Nanosight NTA 3.3 software (Malvern Panalytical Inc) with a detection threshold of 5. Particle counts were normalized for dilution on the machine, dilution of the final pellet, and starting material for exosome extraction. The average count was then taken for fractions V and VI.

### Atomic force microscopy (AFM)

Ten μL of EVs (∼1 μg/μL) were incubated with 100 μL 0.5% sarkosyl (#61747-100ML, Sigma-Aldrich) for 30 min on ice in ultracentrifuge-compatible Beckman microcentrifuge tubes for solubilization of vesicles, and dfPBS was added to 1.2mL. The sample was ultracentrifuged at 100,000 × *g* for 70min at 4°C. The supernatant was removed but leaving 50 μL, and dfPBS was added to 1.2mL for second ultracentrifugation at 100,000 × *g* for 70 min at 4°C. The pellet was dissociated in 10 μL dfPBS, and subjected to AFM imaging by ScanAsyst mode with Multimode 8 AFM machine (Bruker, Billerica MA) as previously described [63].

### Transmission Electron microscopy (TEM)

TEM of EVs was conducted as previously described [4, 53]. Briefly, 5 μL of the EV sample was adsorbed for 1 min to a carbon-coated grid (# CF400-CU, Electron Microscopy Sciences) that had been made hydrophilic by a 20-sec exposure to a glow discharge (25mA). Excess liquid was removed with a filter paper (#1 Whatman), the grid was then floated briefly on a drop of water (to wash away phosphate or salt), blotted on a filer paper, and then stained with 0.75% uranyl formate (#22451 EMS) for 15 seconds. After removing the excess uranyl formate with a filter paper, the grids were examined in a JEOL 1200EX Transmission electron microscope and images were recorded with an AMT 2k CCD camera.

### ELISA of brain tissue extraction and EV samples

Brain tissue homogenate and EV samples were diluted 1:10 in 8M guanidine buffer so solubilization, followed by dilution in TENT buffer (50 mM Tris HCl pH 7.5, 2 mM EDTA, 150mM NaCl, 1% Triton X-100) supplemented with phosphatase inhibitors (Pierce HALT inhibitor), and subjected to human total tau ELISA (human tau: # KHB0042, Thermo Fisher Scientific) according to manufacturer’s instructions.

### EV labelling with PKH26

EVs were labelled with lipophilic red fluorescent dye (PKH26, Sigma-Aldrich), according to the manufacturer’s protocol. Briefly, 0.32-μL PKH26 dye was mixed with 10 μL EV samples in 40 μL diluent C, and incubated for 5 min at room temperature. dfPBS was used as a negative control. The labelling reaction was stopped by adding 50 μL chilled dfPBS, and subjected to Exosome Spin Columns (MW 3000, ThermoFisher, cat.4484449) at 750 × *g* for 2 min to remove the free dye and enrich the labelled EVs, which was adjusted to 5 μg/100 μL for the neuronal EV uptake assay.

### Primary tissue culture of murine cortical neurons

Primary murine cortical neurons were isolated from E16 embryos from pregnant CD-1 mice (Charles River Laboratory). Dissociated cortical tissues were digested with trypsin-EDTA (diluted to 0.125%, #25200072, Invitrogen), triturated by polished pipettes, and strained into single neurons using a 40-μm pore size Falcon cell strainer (Thermo Fisher Scientific), and plated onto 12-mm #1 thickness coverslips or plates, precoated with 100 μg/mL poly-D-lysine (Sigma-Aldrich) diluted in borate buffer (0.05 M boric acid, pH 8.5) and washed with sterile water prior, at 375,000 cells per coverslip in 24-well plates. Neurons at DIV7 were treated with PKH26-labeled EVs for EV uptake or tau transfer study.

### Tau seeding assay

HEK-TauRD P301S FRET cells were plated at in 96-well PDL coated plate (# 354461, Corning) in growth media (DMEM, 10%FBS). The day after, human brain-derived EVs were mixed with 80 μL Opti-MEM and 20 μL Lipofectamine 2000, and incubated at room temperature for 10 min. Subsequently, growth media was removed from the cells, replaced with samples containing Lipofectamine, and incubated at 37°C, 5%CO_2_. After 1 h, Lipofectamine-containing media was removed from the cells and replaced with growth media. Cells were maintained in culture at 37°C, 5% CO_2_ for 72 h afterward. The day of the analysis, cells were washed in PBS, detached with Trypsin 0.25% (#25200072, Invitrogen) and washed with FACS buffer (PBS + 0.5% BSA). Subsequently, cells were fixed in 2%PFA, 2% Sucrose for 15 min at 4 °C, spun at 12,000 rpm for 15 min at 4 °C, resuspended in FACS buffer and acquired with a 5 lasers system LSRII (Becton Dickinson), using pacific-orange and pacific-blue dyes for YFP and CFP, respectively. Data was analyzed by FlowJo and expressed as Integrated FRET Density.

### Stereotaxic surgery

B6 mice at 18–19 months old were deeply anesthetized with isoflurane and immobilized in a stereotaxic frame (David Kopf Instruments) installed with robot stereotaxic injection system (Neurostar). Animals were unilaterally inoculated with human brain-derived EVs or tau aggregates in the dorsal hippocampal OML (bregma: −2.18 mm; lateral: 1.13 mm; depth: −1.9 mm from the skull) using a 10-μL Hamilton syringe as previously described [4]. Each injection site received 1.0 μL of inoculum, containing 300 pg tau /μL for EV samples, and 300 pg or 2 μg of tau per μL oligomeric and fibril fractions.. We noted that majority of the injected materials were deposited at the OML of the hippocampus (Fig. 3A).

### Immunochemistry and Immunofluorescence

Brains were removed after transcardial perfusion fixation with ice-cold 4% paraformaldehyde/PBS followed by post-fixation for 16h and cryoprotection with 15% then 30% sucrose/PBS over 3-5 days. They were cut coronally in 20-µm thickness using a cryostat, and three hippocampal sections separated at least 200 µm per mouse per antibody were used for IHC. The sections were processed by antigen retrieval with Tris-EDTA (pH 8.0) at 80°C, permeabilized in 0.5% Triton-X 100/PBS, and blocked in 10% normal goat serum, 1% BSA, and 0.1% tween-20 in PBS. Sections were incubated GAD67 (#PA5-36054, ThermoFisher scientific); GAD67-biotin-conjugated (# MAB5406B, Millipore), AT8 (# MN1020, ThermoFisher scientific), GluR2/3 (# AB1506, Millipore), MAP-2 (# mab3418, Millipore sigma), c-fos (# 226 003, Synaptic Systems), PS422 (# 44-764G, ThermoFisher scientific), Alz50 and MC1, CP13, PHF-1 (as kind gifts provided by Dr. Davis Peter), diluted with 1% BSA, 0.025% tween-20 in PBS at 4°C for overnight (see Supplementary Table S3 for antibody information). Sections were then washed and incubated in secondary antibodies (AlexaFluor 647 goat anti-mouse; 1:1000, AlexaFluor488 goat anti-rabbit; 1:1000, AlexaFluor568 streptavidin 1:1000) for 1 h at room temperature. All images were captured on Nikon deconvolution wide-field epifluorescence system (Nikon Instruments) or confocal microscopic imaging as described below.

### Confocal image processing and quantification by Imaris

All confocal imaging was performed on a LSM710 using Zen 2010 software (Zeiss) or a Leica TCS SP8 lightning microscope at the inverted Leica DMi8 microscope stand using the confocal mode with a 63× oil immersion/1.4 N.A objective using a 1.1 optical zoom at a pinhole of 1.0 Airy units. Images of 2048 × 2048 pixels as confocal stacks with a z-interval of 0.28 μm system optimized was used to image cells. For imaging GAD67 puncta, a 552-nm laser line was used and emission was collected at 565–650 nm; for imaging c-fos, a 488-nm laser line was used and emission was collected at 490-600 nm. Gain and off-set were set at values which prevented saturated and empty pixels. After image acquisition, all images were applied with lightning deconvolution. The quantification of GAD67 positive puncta was counted by using the “spot” module of Imaris 9.5, 64-bit version (Bitplane AG, Saint Paul, MN, www.bitplane.com). Manual cutting of the CA1 pyramidal cells with GAD67 fields in 3D. This program analyzes stacks of confocal sections acquired in two channels (red for GAD67, blue for DAPI represents cell number). Final data analysis was performed using Microsoft Excel and Graph rendering was done in GraphPad Prism

### Biochemical sequential extraction from mouse brains

Brain tissues were removed from CTRL EV, pAD EV and AD EV-injected mice at the designated time points after transcardial perfusion of animals by ice-cold PBS to minimize contamination of blood-derived mouse immunoglobulins. Hippocampal and cortical regions were dissected separately, snap frozen in dry ice and stored at −80 °C before protein extraction. For enrichment of tau oligomers and fibrils, sequential extractions were performed as follows: Each hippocampal tissue was homogenized in 9 volumes of TBS buffer (50 mM Tris-Cl, pH 8.0 in saline) supplemented with protease and phosphatase inhibitor cocktails (# PI78443, Thermo Fisher Scientific). The homogenate was centrifuged at 48,300 × *g* for 20 min at 4 °C. The supernatant and pellet are designated as S1 (TBS-supernatant) and P1 (TBS-pellet) fraction, respectively. The S1 fraction was ultracentrifuged at 186,340 × *g* at 4 °C for 40 min. The pellet fraction (S1p) was resuspended in a 4 volume of double-filtered TE buffer relative to the starting weight of the tissue, aliquoted and frozen at −80°C as tau oligomer-enriched fraction. The P1 fraction was resuspended in 5 volume of wet weight of the original tissue of buffer B (1% sarkosyl, 10 mM Tris, pH 7.4, 800 mM NaCl, 10% sucrose, 1 mM EGTA, 1 mM PMSF, all from Sigma-Aldrich) and incubated by rotating with the bench top thermomixer at 37 °C for 1 h. The sample was ultracentrifuged at 186,340 × *g* for 1 h at 4 °C. After completely removing the supernatant and rinsing the pellet in sterile PBS, sarkosyl-insoluble pellet (P3) was resuspended with 50 μL double-filtered TE buffer (10 mM Tris, 1 mM EDTA, pH 8.0), aliquoted and frozen at −80°C as tau fibril-enriched fraction.

### Western and dot blotting

For western blotting, homogenates (Ho) of hippocampus from each experimental group and an equal proportion of corresponding Ho, S1, S1p and P3, were loaded on 10% SDS–PAGE gels (Bio-Rad) and electro-transferred to 0.45-μm nitrocellulose membranes (Bio-Rad). For dot blotting, an equal volume of EVs sample were dotted onto 0.45-μm nitrocellulose membranes (Bio-Rad) and washed twice with TBS buffer. The nitrocellulose membranes were then blocked in freshly prepared 5% skim milk diluted in TBS before being immunoblotted with specific primary antibodies (Supplementary Table 3). The membrane was further incubated with HRP-labeled secondary antibodies and scanned using C300 digital chemiluminescent imager (Azure Biosystems). The optical densities were measured using Image J software.

### Whole-cell patch clamp recording

#### Preparation of Brain Slices for Recording and Filling

Immediately after decapitation, mouse brains were rapidly removed and placed in oxygenated (95% O_2_ and 5% CO_2_) ice-cold Ringer’s solution containing following ingredients (in mM): 25 NaHCO_3_, 124 NaCl, 1 KCl, 2 KH_2_PO_4_, 10 glucose, 2.5 CaCl_2_, 1.3 MgCl_2_ (pH 7.4; Sigma-Aldrich). A total of four to five 300-µm thick acute coronal sections containing the hippocampus were obtained from each subject. Over an 8-10 h period, slices were individually transferred from the incubation chamber to submersion-type recording chambers (Harvard Apparatus, Holliston, MA) affixed to the stages of Nikon E600 infrared-differential interference contrast (IR-DIC) microscopes (Micro Video Instruments, Avon, MA) with a water-immersion lens (40×, 0.9 NA; Olympus) for recording. During recordings, slices were superfused in room-temperature Ringer’s solution bubbled with carbogen (95% O_2_, 5% CO_2_) a rate of 2.5 ml/min. Whole-cell patch clamp recordings were obtained from the soma of visually identified CA1 pyramidal cells in both the dorsal and ventral hippocampus of ipsilateral side of the brain. Electrodes were created from borosilicate glass with a Flaming and Brown micropipette puller (Model P-87, Sutter Instruments). These pulled patch pipettes were filled with potassium methanesulfonate (KMS) based intracellular solution, with concentrations in mM as follows: (KCH_3_SO_3_ 122, MgCl_2_ 2, EGTA 5, Na-HEPES 10, Na_2_ATP 5)., and had a resistance of 5.5–6.5 MΩ in external Ringer’s solution.

#### Physiological Inclusion Criteria

Single AP properties (including threshold, amplitude, Action potential Half-Width (APHW), rise and fall) were measured on the second evoked AP in a 200 ms current-clamp series that preferentially evoked 3 or more action potentials after depolarizing step-current. We proceeded to High Rn or Low Rn only if neurons were unable to elicit AP at 200 ms. AP half-width was computed at half-max of AP amplitude, where the amplitude was measured from the threshold to the absolute peak of the spike. All the quantification for AP properties was carried out in an expanded timescale, and the linear measure tool we used in FitMaster analysis software (HEKA Elektronik) to measure all single AP properties. An algorithm designed in Matlab was used to automatically detect these parameters. In the few cases where it failed to do so, a manual detection method was used. The final paradigm in the Current-clamp configuration was to inject 2 s hyperpolarizing and depolarizing steps (−100 to +120 pA with increments of 20 pA or −220 pA to +330 pA with increments of 50pA, 12.5kHz sampling frequency) to assess repetitive AP firing. Those neurons which did not fire repetitively in depolarizing step were discarded. Firing rates in response to current steps were analyzed fitting with a generalized linear model, using the genotype, CA1 pyramidal cells types, rheobase, input resistance, injected current level and their respective interactions as independent variables. Whole-cell voltage clamp was used to measure AMPA receptor-mediated spontaneous excitatory currents (sEPSCs) response for 2 min at a holding potential of −80 mV (6.67 kHz sampling frequency). The same neuron was held at −40 mV (6.67 kHz sampling frequency) for 2 min to obtain enough sample size to measure GABA receptor-mediated spontaneous inhibitory currents (sIPSCs). All recorded traces were run through Minianalysis software (Synaptosoft) which allowed for quantification of synaptic current properties such as frequency, amplitude, area, time to rise and time to decay. To determine the kinetics of EPSCs and IPSCs, the rise and decay of averaged traces were each fit to a single-exponential function. In all of the synaptic current measurements, the event detection threshold was set at the maximum root mean squared noise level (5 pA). All neurons had resting membrane potentials between −55 and −75 mV (somatic recordings) and were confirmed to have intact somas and apical tufts.

## Statistical Analyses

All data are presented as means ± standard error of the mean (s.e.m). Comparisons between two groups were done by two-tailed paired or unpaired Student’s *t*-tests. Multiple comparisons were performed by either one- or two-way ANOVA, followed by Turkey’s or Bonferroni’s *post hoc*. Statistical analyses were performed using Prism 8.0 (GraphPad Software). A statistically significant difference was assumed at p <0.05.

## Acknowledgement

We would like to thank JC Delpech, A DeLeo, and other members of the Laboratory of Molecular NeuroTherapeutics for scientific suggestions and technical assistance, and Dr. Peter Davies for the generous gift of the Alz50, MC1, CP13 and PHF1 monoclonal antibodies.

## Author contributions

Conceptualization, Z.R., A.M.D., S.I., J.L. and T.I.; Methodology, Z.R., D.P., A.Y-K., S.M., S-V.K., K.T-K., S.G., R.K., H.E.G., Y.W., J.L.; Bioinformatics: J.H.; Image analysis, Z.R, J.H, A.Y-K., S-V.K., YW; Manuscript writing & Editing, Z.R., D.P., T.I., S.I.; All authors read and approved the final manuscript.

## Figure legends

**Supplementary Figure S1. Dot blot of Tau KO and human brain-derived EV samples by tau oligomer-specific antibodies.**

**Supplementary Figure S2. AT8 staining of young and aged mice after the injection of human brain-derived EVs.** Young (2 months old) or Aged (18 months old) mice were stereotaxically injected with human brain-derived EVs (CTRL, pAD or AD) containing 300 pg tau in 1 μL volume into the outer molecular layer of dentate gyrus, and sacrificed 4.5 months after the injection for the neuropathological examination using AT8 (pSer202/pSer205 tau) monoclonal (red) and counterstained with Dapi (blue). Coronal sections depicting the ipsilateral hippocampal region (left) and hilus region (right). Sale bars: 200 μm (left) and 50 μm (right)

**Supplementary Figure S3. Tau pathology staining with Alz50, MC1, CP13, PS422 and PHF1 antibodies.** Representative images of AT8 (pSer202/pSer205 tau, A), Alz50 (conformation-specific misfolded tau, B), MC1 (conformation-specific misfolded tau, C), CP13 (pSer202 tau, D), PS422 (pSer422 tau, E) and PHF1 staining (pSer396/pSer404 tau, F) (red) and Dapi (blue) 4.5 months after intrahippocampal injection of saline, Tau KO EV, CTRL EV, and pAD EV or AD EV (1 μL volume containing 300 pg tau) into aged B6 mouse brain. Scale bar = 200 (left) and 100μm (right).

**Supplementary Figure S4. Tau accumulation in the injection site of cortex 4.5 months post intracranial injection.** Aged mice (18 months of age) were intracranially injected with AD EV (left), tau oligomer-enriched fraction (middle) or tau fibril-enriched fraction (right) containing 300 pg tau in 1 μL volume. The animals were sacrificed and tested for neuropathology using AT8 (red) and counterstained by Dapi (blue) for nuclear staining. Perikaryal accumulation of p-tau in AD EV-injected cortical region (left) and neuropil staining in tau oligomer (middle) or fibril-injected cortical region (right). Scale bar=50 μm.

**Supplementary Figure S5. Injection of 300pg of EV-tau, 2 μg of oligomeric or fibril tau derived from AD brain induced tau propagation in mouse brain**

**a.** Representative images of AT8 immunostained recipient mice after unilateral injection of 300 pg of AD EVs (left), 2 μg of tau oligomer-enriched fraction (middle) and tau fibril-enriched fraction (right) in cortical region (top panels, scale bar=200 µm) and dentate gyrus (bottom panels, scale bar=50 µm).

**b.** Quantification of AT8^+^ neurons in the hippocampus of recipient mice. *n.s* denotes no significance as determined by one-way ANOVA (alpha = 0.05) and Turkey’s *post-hoc*. EV-tau, oligomeric and fibril tau group: n = 3-6 mice per group for quantification. Bregma −1.34 to −3.64, 4 sections per mouse were analyzed. Each dot represents mean value per animal. Graphs indicate mean ± s.e.m.

**c.** Representative images of PHF1 immunoblotting of the same injected amount of isolated EVs, tau oligos and tau fibrils by PHF1 antibodies.

**Supplementary Figure S6. AT8**^**+**^ **cell was co-stained with parvalbumin**^**+**^ **inhibitory but not neurogranin**^**+**^ **excitatory neurons in the hippocampal region after AD EV injection.**

**a.** AT8 (pSer202/pSer205 tau, red) and parvalbumin (inhibitory neuronal marker, green) immunostaining in the ipsilateral CA1, CA3 and DG regions of hippocampus from AD EV injected mice at 4.5 months post injection. Nuclei were counterstained by Dapi (blue). Scale bar= 50 μm.

**b.** AT8 (red) and neurogranin (excitatory neuronal marker, green) immunostaining of the same brains. Nuclei were counterstained by Dapi (blue). Scale bar= 50 μm.

**Supplementary Table S1. Demographics of human cases used in the study**

**Supplementary Table S2. Biochemical characterization of EV-enriched fractions derived from human brain**

**Supplementary Table S3. List of antibodies used in the study**

**Supplementary Table S4-S8. Electrophysiological properties of CA1 pyramidal cells in brain-derived EV-injected mouse brain**

