## Supplementary Figures S1-S6 for "Alzheimer’s disease brain-derived tau-containing extracellular vesicles: Pathobiology and GABAergic neuronal transmission"

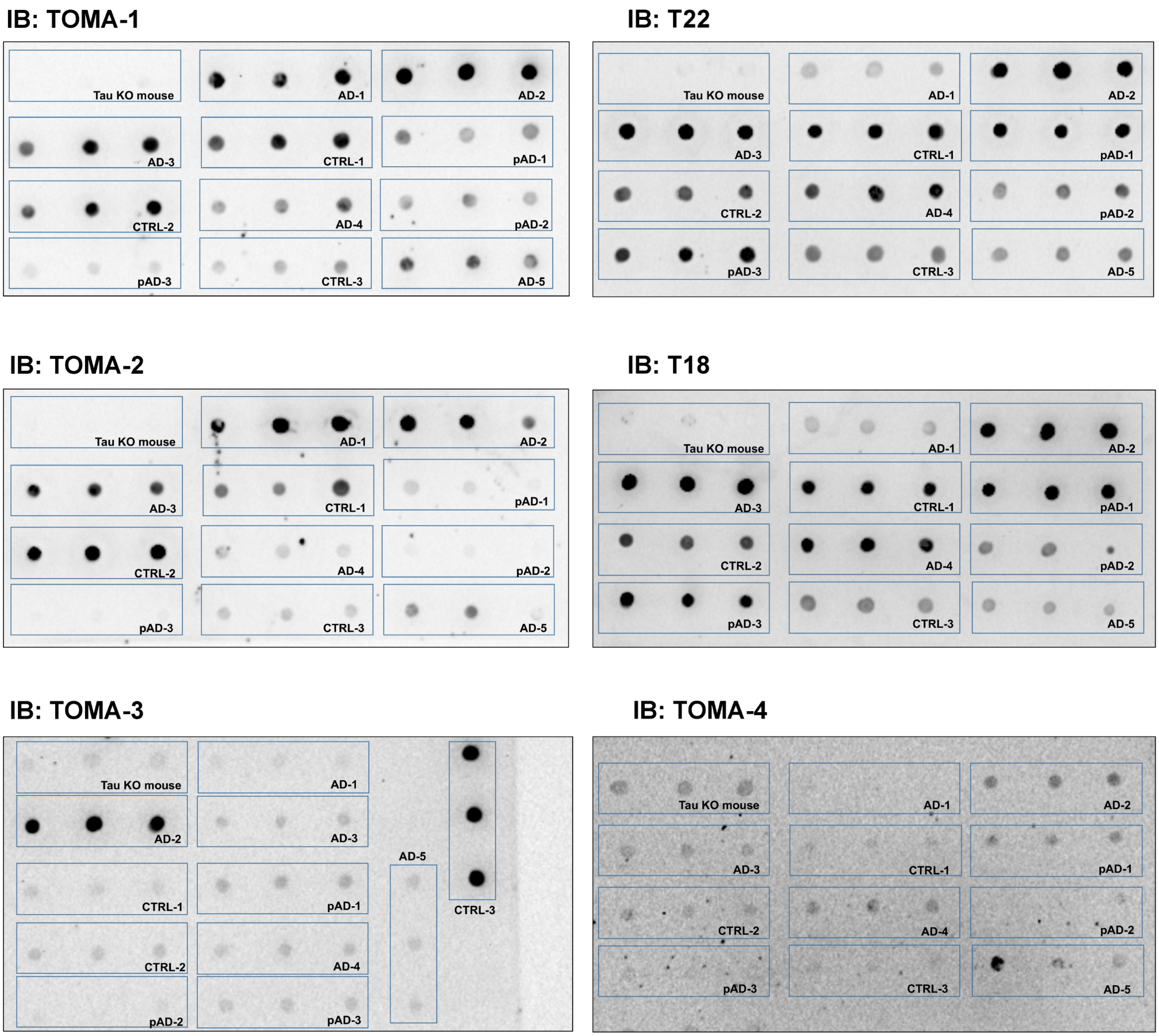


**Supplementary Figure S1. Dot blot of Tau KO and human brain-derived EV samples by tau oligomer-specific antibodies.**

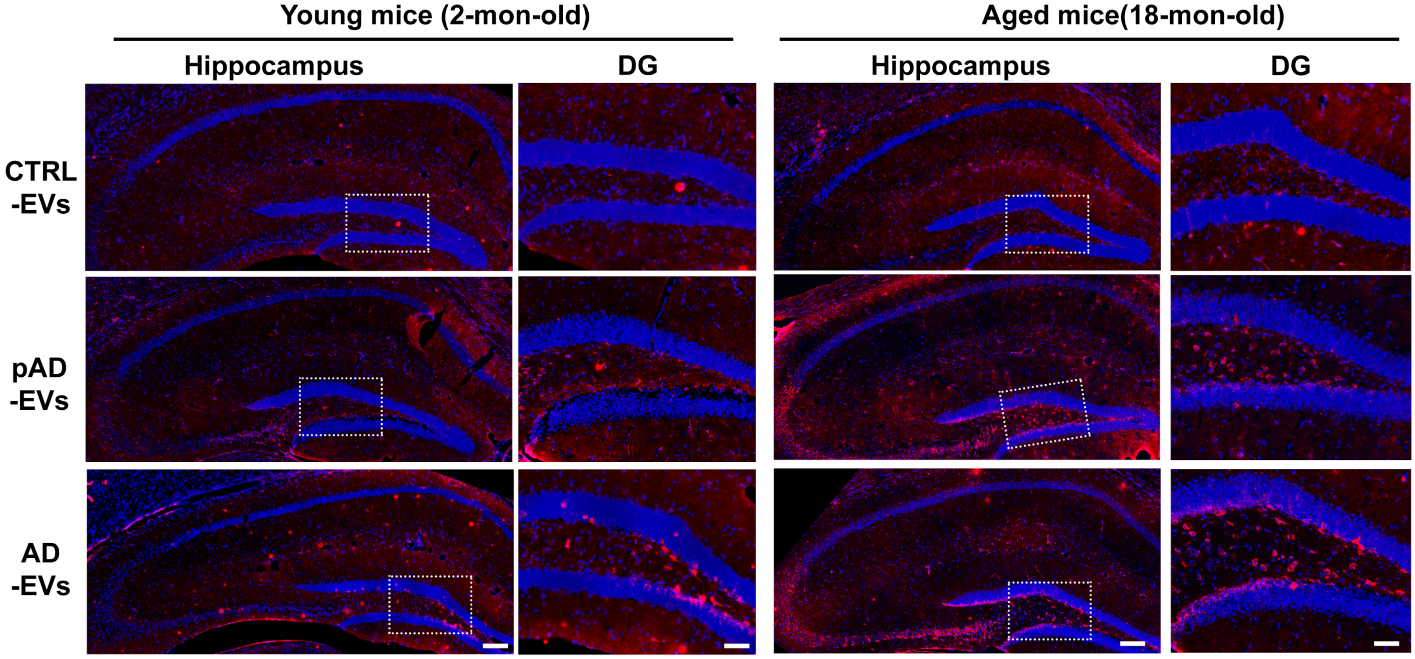


**Supplementary Figure S2. AT8 staining of young and aged mice after the injection of human brain-derived EVs.** Young (2 months old) or Aged (18 months old) mice were stereotaxically injected with human brain-derived EVs (CTRL, pAD or AD) containing 300 pg tau in 1 µL volume into the outer molecular layer of dentate gyrus, and sacrificed 4.5 months after the injection for the neuropathological examination using AT8 (pSer202/pSer205 tau) monoclonal (red) and counterstained with Dapi (blue). Coronal sections depicting the ipsilateral hippocampal region (left) and hilus region (right). Sale bars: 200 µm (left) and 50 µm (right)

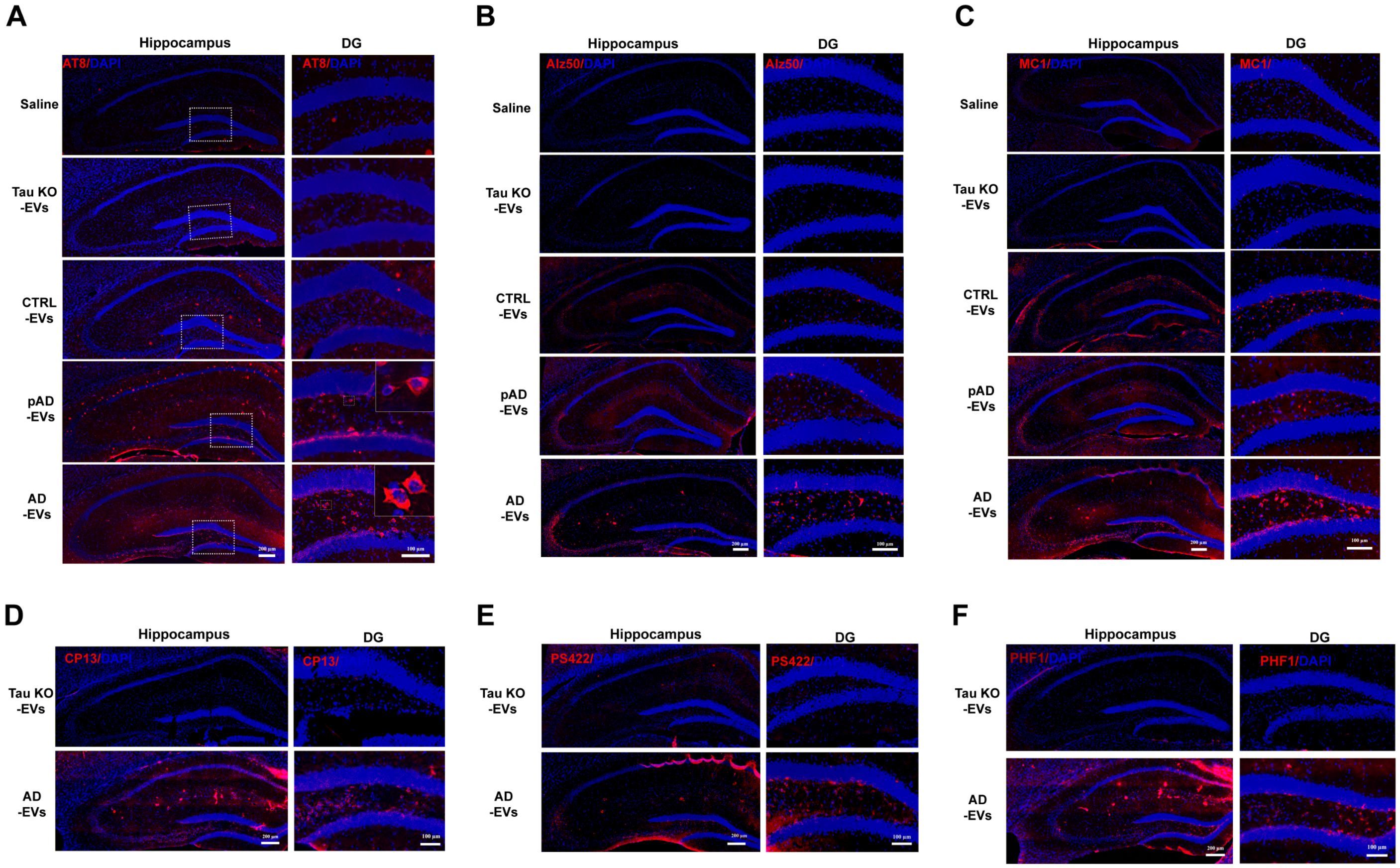


**Supplementary Figure S3. Tau pathology staining with Alz50, MC1, CP13, PS422 and PHF1 antibodies.** Representative images of AT8 (pSer202/pSer205 tau, A), Alz50 (conformation-specific misfolded tau, B), MC1 (conformation-specific misfolded tau, C), CP13 (pSer202 tau, D), PS422 (pSer422 tau, E) and PHF1 staining (pSer396/pSer404 tau, F) (red) and Dapi (blue) 4.5 months after intrahippocampal injection of saline, Tau KO EV, CTRL EV, and pAD EV or AD EV (1 μL volume containing 300 pg tau) into aged B6 mouse brain. Scale bar = 200 (left) and 100μm (right).

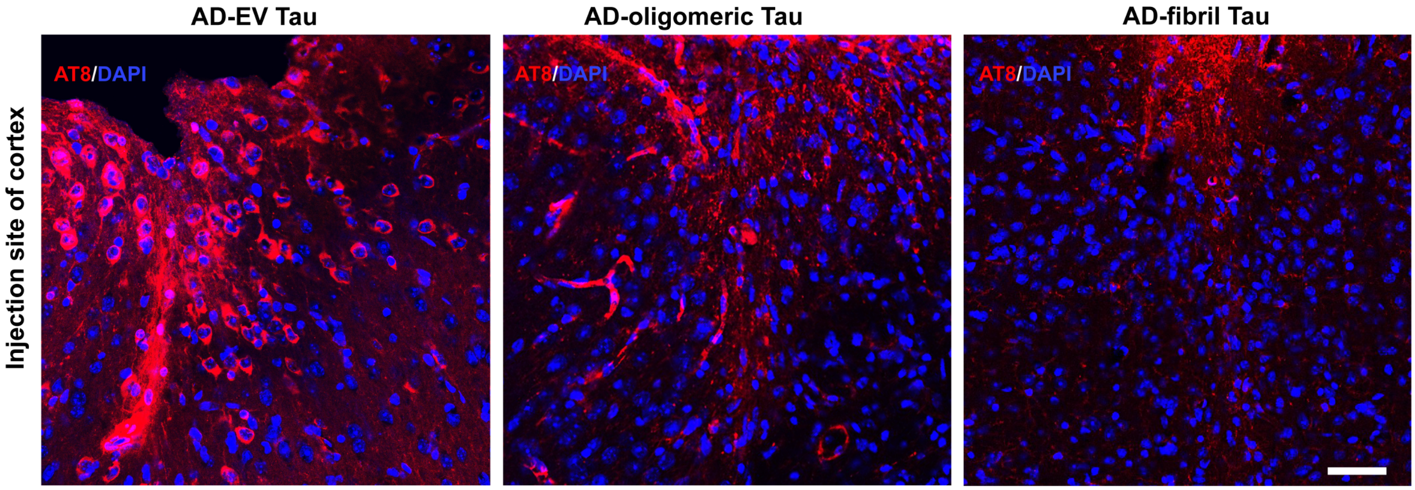


**Supplementary Figure S4. Tau accumulation in the injection site of cortex 4.5 months post intracranial injection.** Aged mice (18 months of age) were intracranially injected with AD EV (left), tau oligomer-enriched fraction (middle) or tau fibril-enriched fraction (right) containing 300 pg tau in 1 μL volume. The animals were sacrificed and tested for neuropathology using AT8 (red) and counterstained by Dapi (blue) for nuclear staining. Perikaryal accumulation of p-tau in AD EV-injected cortical region (left) and neuropil staining in tau oligomer (middle) or fibril-injected cortical region (right). Scale bar=50 μm.

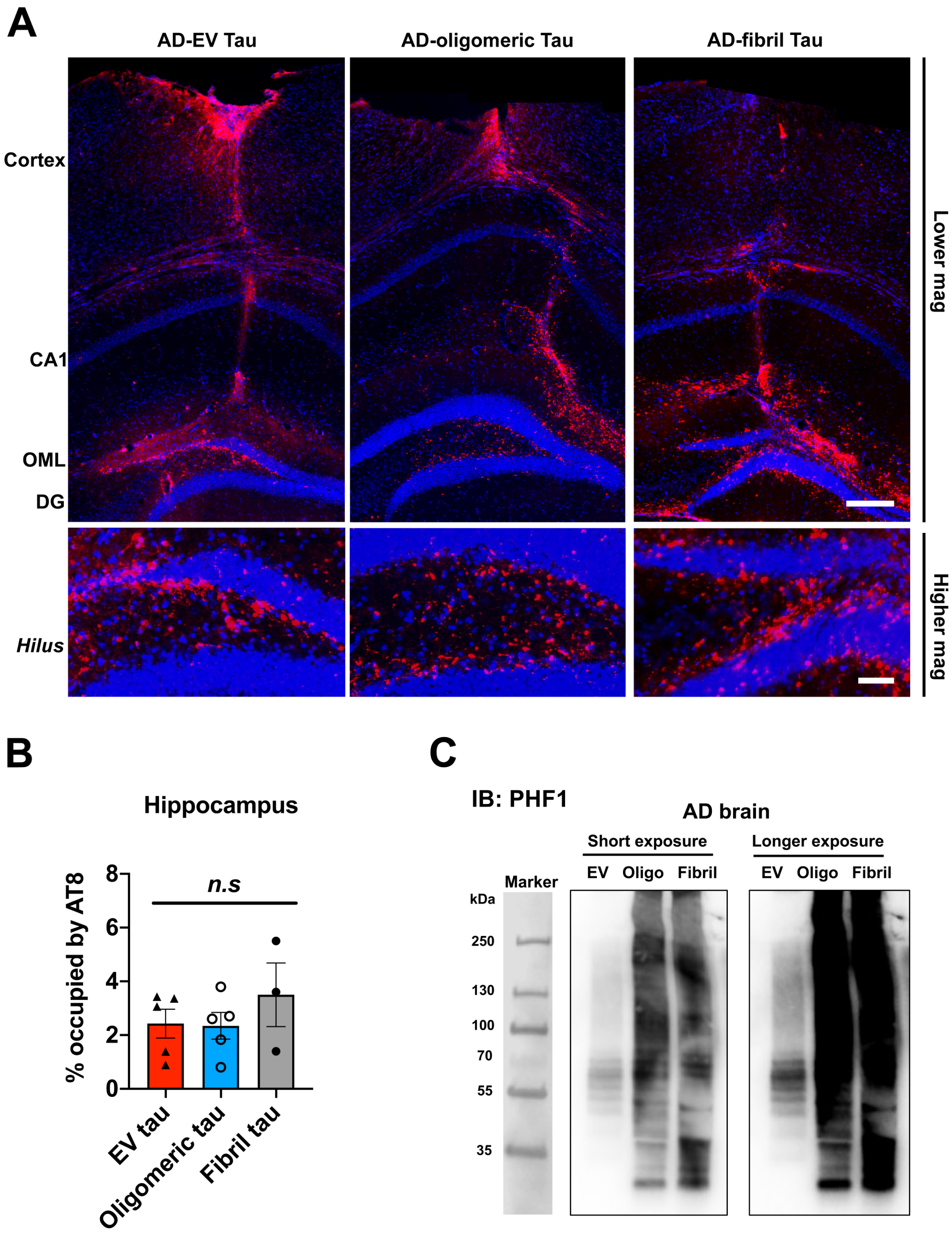


**Supplementary Figure S5. Injection of 300 pg of EV-tau, 2 µg of oligomeric or fibril tau derived from AD brain induced tau propagation in mouse brain**

**a.** Representative images of AT8 immunostained recipient mice after unilateral injection of 300 pg of AD EVs (left), 2 µg of tau oligomer-enriched fraction (middle) and tau fibril-enriched fraction (right) in cortical region (top panels, scale bar=200 µm) and dentate gyrus (bottom panels, scale bar=50 µm).

**b.** Quantification of $AT8^{+}$ neurons in the hippocampus of recipient mice. *n.s* denotes no significance as determined by one-way ANOVA (alpha = 0.05) and Turkey's *post-hoc*. EV-tau, oligomeric and fibril tau group: n = 3-6 mice per group for quantification. Bregma -1.34 to -3.64, 4 sections per mouse were analyzed. Each dot represents mean value per animal. Graphs indicate mean ± s.e.m.

**c**. Representative images of PHF1 immunoblotting of the same injected amount of isolated EVs, tau oligos and tau fibrils by PHF1 antibodies.

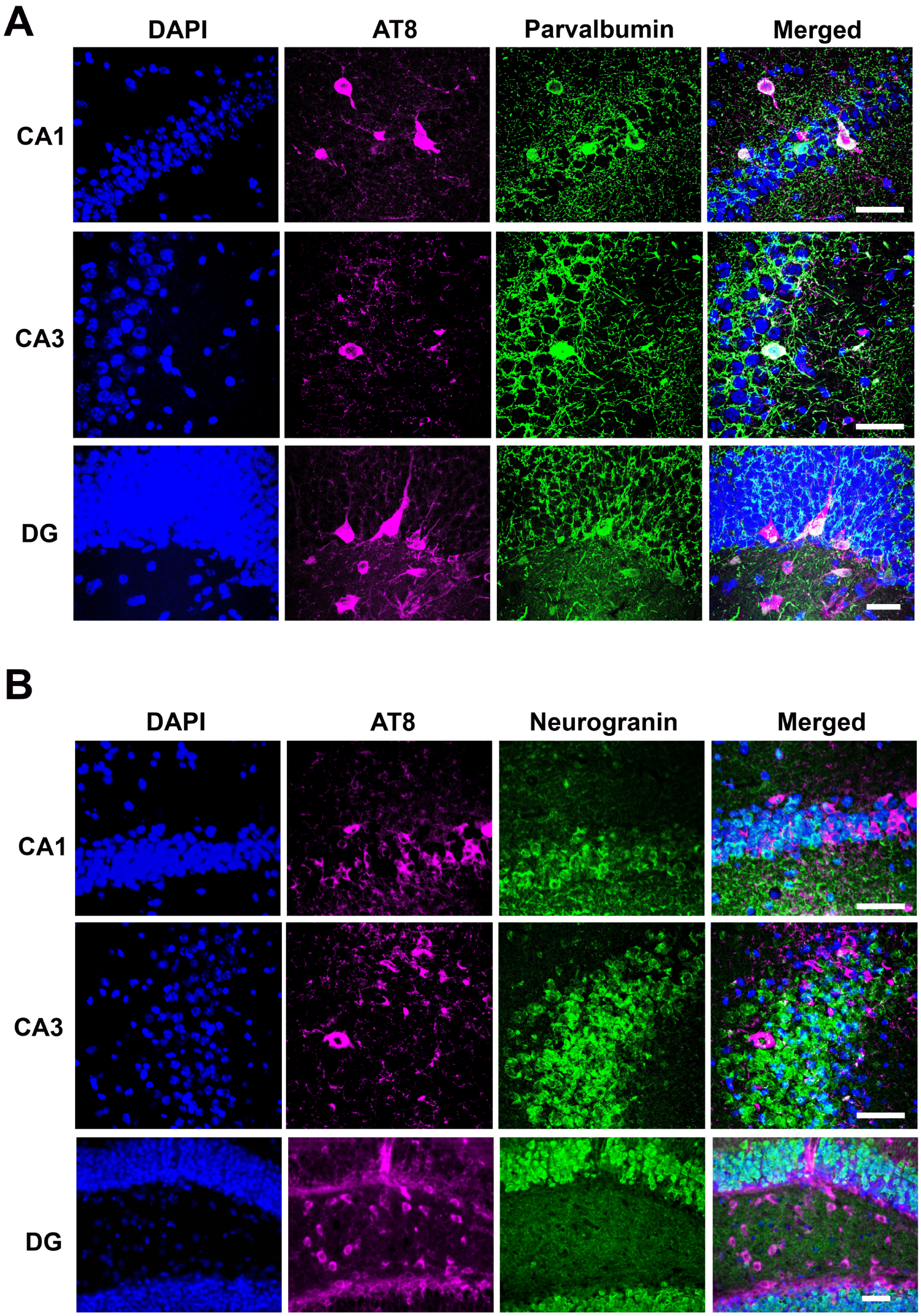


**Supplementary Figure S6. AT8$^+$ cell was co-stained with parvalbumin$^+$ inhibitory but not neurogranin$^+$ excitatory neurons in the hippocampal region after AD EV injection.**

**a**. AT8 (pSer202/pSer205 tau, red) and parvalbumin (inhibitory neuronal marker, green) immunostaining in the ipsilateral CA1, CA3 and DG regions of hippocampus from AD EV injected mice at 4.5 months post injection. Nuclei were counterstained by Dapi (blue). Scale bar= 50 µm.

**b.** AT8 (red) and neurogranin (excitatory neuronal marker, green) immunostaining of the same brains. Nuclei were counterstained by Dapi (blue). Scale bar= 50 µm.
